# EGF signaling in bowel carcinoma cells utilizes higher order architectures of EGFR and HER2

**DOI:** 10.1101/2020.07.11.198572

**Authors:** Adam J. M. Wollman, Charlotte Fournier, Isabel Llorente-Garcia, Oliver Harriman, Alex L. Hargreaves, Sviatlana Shashkova, Peng Zhou, Ta-Chun Liu, Djamila Ouaret, Jenny Wilding, Akihiro Kusumi, Walter Bodmer, Mark C. Leake

**Author notes:** These authors contributed jointly to this work (Adam J. M. Wollman, Charlotte Fournier, Isabel Llorente-Garcia, Oliver Harriman). Correspondence should be addressed to M.C.L.

## Abstract

Epidermal growth factor (EGF) signaling regulates normal cell development, however EGF receptor (EGFR) overexpression is reported in several carcinomas. Despite structural and biochemical evidence that EGF-EGFR ligation activates signaling through monomer-dimer transitions, live cell mechanistic details remain contentious. We report single-molecule multispectral TIRF of human epithelial carcinoma cells transfected with fluorescent EGFR, and of CHO-K1 cells containing fluorescent EGFR and HER2, enabling super-resolved localization to quantify receptor architectures and spatiotemporal dynamics upon EGF ligation. Using inhibitors that block binding to EGFR, and time-dependent kinetics modelling, we find that pre-activated EGFR consist predominantly of preformed clusters that contain a mixture of EGFR and HER2, whose stoichiometry increases following EGF activation. Although complicated by EGFR internalization and recycling, our observation of an EGFR:EGF stoichiometry >1 for plasma membrane colocalized EGFR/EGF foci soon after activation may indicate preferential binding of EGF ligand to EGFR monomers, negative cooperativity and preferential ligated-unligated dimerization of monomers.

## Introduction

Epidermal growth factor receptor (EGFR) is essential for epithelial tissues and several signaling pathways, its upregulation is implicated in several carcinomas(1). Human EGFR or ERBB1, (‘ErB1’or ‘HER1’) is a protein of receptor tyrosine kinase (RTK) family with three other ERBB members, ERBB2 (‘ErbB2’ or ‘HER2’), ERBB3 (‘ErbB3’ or ‘HER3’) and ERBB4 (‘ErbB4’ or ‘HER4’), expressed in plasma membranes of epithelial cells(2). EGFR has an extracellular region, with subdomains I-IV of which I and III participate in ligand binding(3), connected to a cytoplasmic domain containing a tyrosine kinase.

EGFR activation requires ligand binding, receptor-receptor interactions, and tyrosine kinase activity with 11 different ligands binding to ERBB proteins, including EGF which binds to EGFR(4). Subsequent autophosphorylation of intracellular residues initiate reactions stimulating cell growth, differentiation and proliferation, terminated by internalization and proteolytic degradation of receptor-ligand(5).

Much is known about interactions that contribute to signal transduction, however, controversy remains concerning *in vivo* EGFR composition before and after activation and the roles of higher order complexes. Small angle X-ray scattering and isothermal titration calorimetry to EGFR’s isolated extracellular domain (sEGFR) suggest EGF binds to sEGFR monomers, receptor dimerization involving association of two monomeric EGF-sEGFR(6). Multi-angle laser light scattering suggests sEGFR is monomeric in solution but dimeric after EGF ligation(7). Fluorescence anisotropy indicates 1:1 binding of EGF:sEGFR, analytical ultracentrifugation suggesting 2(EGF-sEGFR) complexes(8). Structural evidence indicates activation is preceded by ligand binding to receptor monomers such that EGF induces conformational change by removing interactions that autoinhibit dimerization(9) (positive cooperativity). However, binding studies of full length receptors suggest reduced affinity for subsequent binding (negative cooperativity) mediated through an intracellular juxta-membrane domain(10). It has been shown that EGFR dimers with a single bound EGF can be phosphorylated(11). A prediction from negative cooperativity is that EGFR:EGF complexes have a nominal relative stoichiometry of 2:1(12).

Similarly, the first single-molecule fluorescence imaging studies in cells suggested binding of one EGF to a preformed EGFR dimer, rapidly followed by a second to form a 2:2 complex(13). Förster resonance energy transfer (FRET) suggest preformed oligomeric EGFR(14) supported by autocorrelation analysis(15), bimolecular fluorescence complementation (BiFC)(16), and pixel brightness analysis of GFP-labelled EGFR(17). Recent light microscopy advances have yielded new insights in conformational changes in EGFR rotation(18). More recent single-molecule analyses of GFP-labelled CHO cells suggest EGFR forms oligomers prior to EGF binding, triggered at physiological EGF levels(19), contrasting with findings in live *Xenopus* oocytes that report mostly monomeric EGFR before activation(20). EGFR clustering is nuanced since it may involve cooperativity not only between EGFR subunits but also other ERBB proteins(16). EGFR’s oligomeric state before and after activation under physiological conditions remains an open question due to limitations in obtaining simultaneous data on stoichiometries of interacting receptors and ligands, dependence of EGF expression on EGFR oligomerization, the presence of fluorescently labelled and dark EGFR, and species-specific cell line differences.

We investigated a human epithelial carcinoma cell line, with negligible native EGFR protein(21), to improve our understanding of EGF binding to EGFR in cancer. We use single-molecule total internal reflection fluorescence (TIRF) on live human colorectal carcinoma cells into which GFP-labelled EGFR had been stably transfected, coupled to nanoscale tracking of tetramethylrhodamine (TMR) stoichiometrically conjugated to EGF (fig. 1A) both in the presence and absence of popular immunotherapy antibodies which inhibit EGF signaling. We find EGFR forms oligomeric clusters prior to EGF binding with a peak stoichiometry of 6. After EGF ligation, measurements of cluster mobility in the presence of inhibitors which target HER2 suggest they contain clusters of both EGFR and HER2, consistent with subsequent TIRF on a dual-label CHO-K1 cell line which shows EGFR and HER clusters interact transiently even before EGF activation with a dwell time of several hundred milliseconds. Following EGF ligation we see evidence for a relative EGFR:EGF stoichiometry greater than 1 (~2:1 considering the ratio of modal averages, ~4:1 from the ratio of mean averages of stoichiometry). Kinetics modelling suggests a combination of preferential binding of ligand to receptor monomers, negative cooperativity for EGFR activation by EGF(22) and preferential ligated-unligated dimerization of monomers.

**Figure 1.**
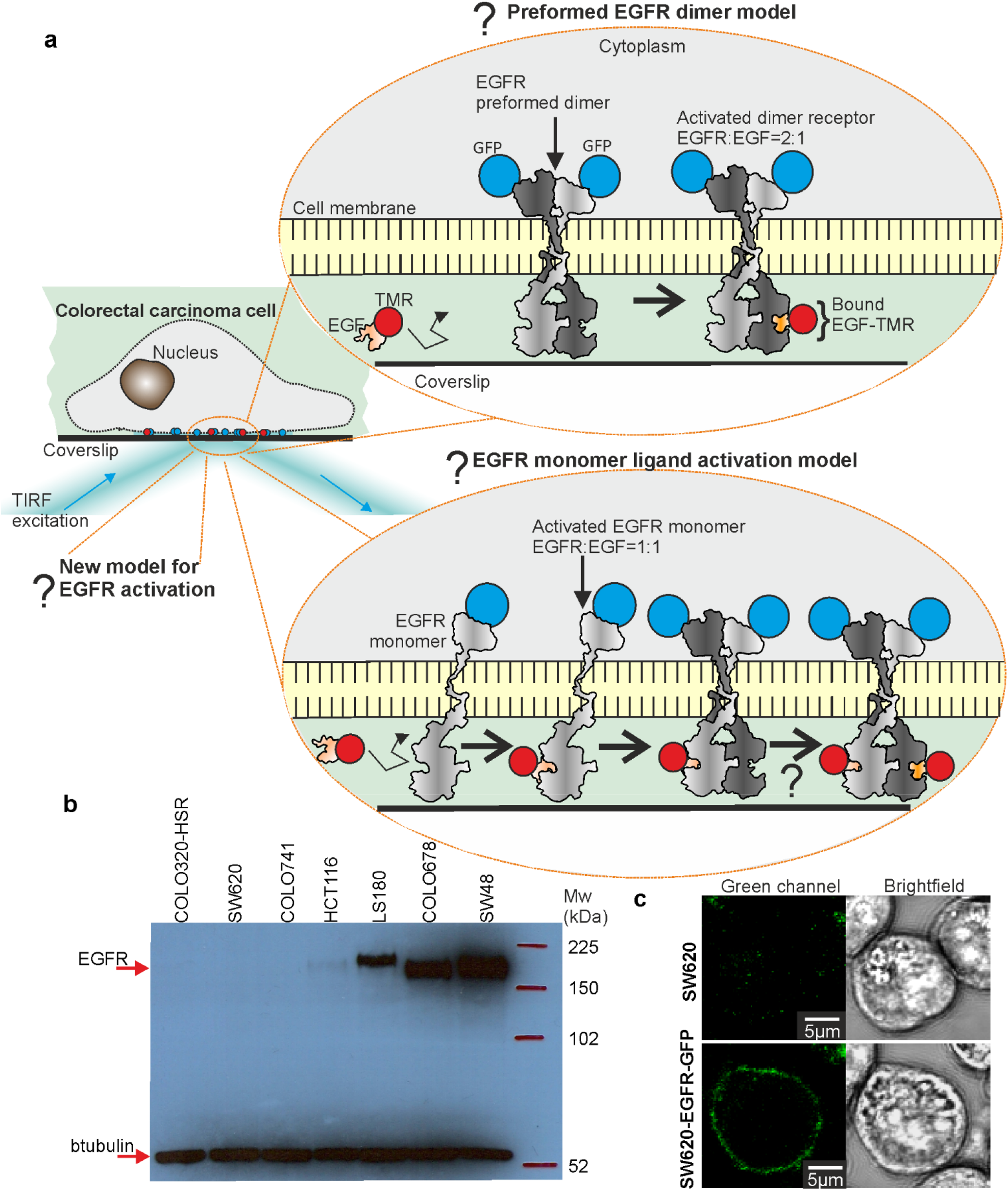
Visualizing EGF-EGFR in human carcinomas. (**A**) TIRF applied to colorectal carcinoma cells. Several models to explain EGFR activation are postulated, including ‘monomer’ and ‘preformed dimer’ models (EGF structure PDB ID 1egf; EGFR monomer and dimer cartoons have been generated by manually combining separate structures with PDB ID values of 1nql, 1ivo, 2jwa, 1m17and 2gs6). (**B**) SDS-PAGE for candidate colorectal carcinoma cell lines, indicating SW620 COLO320-HSR (as opposed to COLO320-DM, its duplicate line) and COLO741 (later found to be a melanoma and not subsequently used) have negligible endogenous EGFR expression compared to positive controls HCT116, LS180, COLO678 and SW48. (**C**) Parental SW620 shows minimal autofluorescence (upper left), while SW620-EGFR-GFP show plasma membrane localization for EGFR-GFP (lower left) from confocal imaging of cells soon after adhering to the coverslip surface focusing at mid-cell-height, fluorescence intensity inside cells comparable to SW620.

## Results

### Construction of EGFR-GFP cells

Human cell line SW620 was selected from a colorectal carcinoma library for low endogenous EGFR expression as quantified by microarray(23) (fig. S1A). EGFR protein was undetectable by western blotting (fig. 1B). SW620 was stably transfected with plasmid pEGFR-EGFP-N1 to give SW620-EGFR-GFP, GFP tagging the cytoplasmic domain far from the EGF binding site. The construct’s kinase activity was confirmed by observing increased phosphorylation of ERK1/2 kinases, EGFR downstream targets, in response to EGF (fig. S1B). RT-qPCR indicated endogenous expression of *HER2*, *HER3* and *HER4* comparable to *EGFR* in the parental SW620, and the construct did not cause significant changes in their expression (fig.1SC). Live cell confocal fluorescence microscopy confirmed plasma membrane localization (fig. S1C), with immunofluorescence on fixed cells using AlexaFluor633-labelled anti-EGFR and anti-GFP antibodies demonstrating colocalization of GFP and EGFR (fig. S2A-D).

### TIRF optimized for single-molecule EGF/EGFR detection

We optimized a bespoke TIRF microscope (fig. S2E) for single-molecule detection using a surface assay(24) in which GFP or EGF-TMR was conjugated to a glass coverslip using IgG/Fab with binding specificity to GFP or EGF (fig. 2A and fig. S3A). After ~1s laser illumination, foci exhibited step-wise photobleaching (fig. 2B), indicative of single GFP or TMR, each with a brightness (summed pixel intensity) of ~2,000 counts on our detector (fig. S3B).

**Figure 2.**
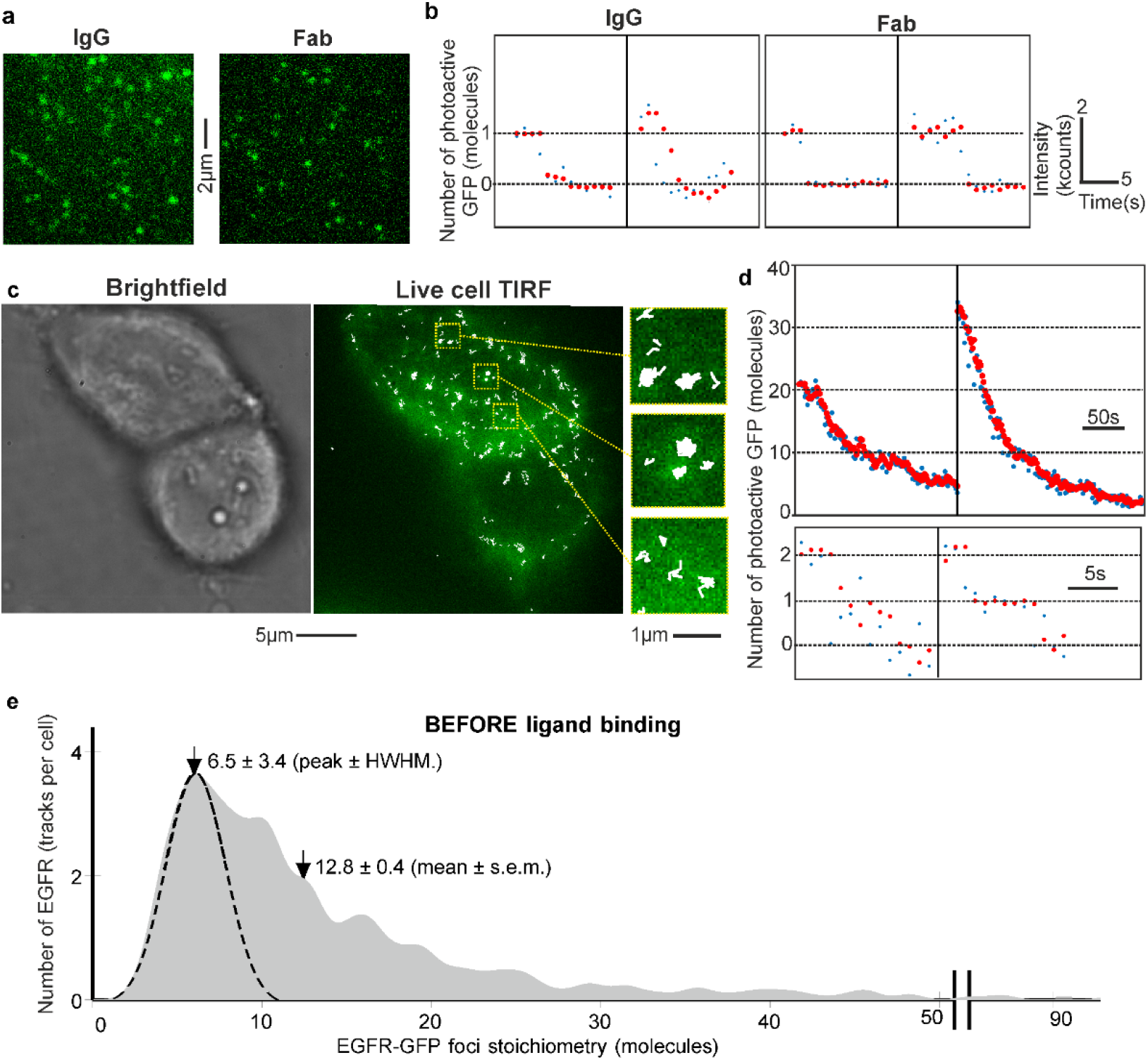
EGFR stoichiometry before EGF binds. **(A)** TIRF of GFP *in vitro* using IgG/Fab conjugation. **(B)** Step-wise photobleaching showing raw (blue) and output data of an edge-preserving filter(28) (red), kilocounts equivalent to counts on our detector x10^3^. **(C)** Two SW620-EGFR-GFP cells showing GFP (green) and overlaid tracking (white) with zoom-ins (inset). **(D)** Photobleaches from tracked EGFR-GFP with stoichiometries of several tens (upper), down to a minimum of two (lower). **(E)** Distribution of EGFR-GFP stoichiometry before EGF ligation showing peak at ~6 and mean 12.8 molecules, N=19 cells, N=1,250 tracks in total (~66 tracks per cell), corresponding to mean of approximately 850 tracked EGFR per cell.

### Tracked EGFR are oligomeric before EGF binds

Before adding EGF in serum-free medium we observed fluorescent foci at a surface density of 0.1-0.4 per μm^2^ in the plasma membrane (fig.2C and fig. S4A). In most cells foci could be detected across the full extent of the basal membrane and exhibited a smooth surface consistent with earlier SEM imaging performed on the SW620 cell line(21), though in a few which exhibited finger-like filopodia protrusions of the membrane we saw a small localization bias towards the cell peripheries. We detected a mean 66±28 (s.d.) foci per cell and monitored their spatiotemporal dynamics over several seconds with 40nm precision using super-resolved tracking(25), indicating both mobile and immobile foci (movie S1). Foci widths were within 10% of those observed for single GFP *in vitro* (~250nm half width at half maximum). However, brightness was greater than that expected for monomeric GFP, exhibiting stochastic photobleaching steps (fig.2D), which we used to determine stoichiometry by dividing the initial brightness by that of single GFP(24). To determine GFP brightness we quantified mean foci brightness towards the end of each photobleach, when only one photoactive molecule remained, indicating live cell values within 15% of that *in vitro* (fig. S3B). Previous live cell measurements using the same fluorescent protein indicate the proportion of immature GFP is <15% of the total(26). We measured a broad range of stoichiometry, across different cells and within the same cell, of 2-90 EGFR molecules per focus, with peak value ~6 and mean 12.8±0.4 molecules (±s.e.m.) (fig. 2E).

We could not detect any monomeric EGFR-GFP before adding EGF, despite our microscope having single GFP sensitivity *in silico* (fig. S4B) and *in vitro* under the same imaging conditions, from >1,000 tracks in 19 different cells. We wondered if random overlap of EGFR-GFP diffraction-limited images which are not physically in the same cluster could account for the apparent stoichiometries of EGFR. We modelled this phenomenon by convolving overlap probability(27) (Methods) with the brightness distribution of a cluster in a range of different oligomeric states from monomers through to tetramers (suggested from a previous single-molecule study(19)), which resulted in very poor agreement to the observed data (fig. S4C *R*^2^<0). However, simulating cluster stoichiometry using a random Poisson distribution whose mean was equal to 6 molecules resulted in reasonable predictions which could account for approximately 50% of the observed variance in the experimental stoichiometry distribution (*R*^2^=0.4923, fig. S4D).

By quantifying summed TIRF pixel intensities for SW620-EGFR-GFP, correcting for autofluorescence and cell area (Methods), we measured the mean EGFR-GFP copy number in the plasma membrane as 200,000±11,000 molecules per cell (±s.e.m.). The 100nm TIRF penetration depth we calculate will illuminate approximately 1/3 of the cell’s plasma membrane (the planar basal membrane in contact with the coverglass plus a portion of the curved membrane away from the surface), so the maximum EGFR-GFP content visible is ~67,000 molecules per cell, whereas what we actually track is ~1% of this. We calculate that the maximum stoichiometry of foci which are not detected as distinct foci is approximately a few tens of molecules (Methods), however, we cannot directly exclude the possibility that monomeric EGFR are present.

### Mean relative stoichiometry of EGFR to ligated EGF is 4:1

To determine the effect of EGF binding on EGFR stoichiometry and spatiotemporal dynamics, live SW620-EGFR-GFP and non-GFP controls were first kept in serum-free media for 24h to minimize binding of serum-based EGFR ligands. We visualized cells using TIRF then added EGF-TMR, enabling simultaneous observation of EGFR and EGF in green/red color channels, before and after EGF ligation. Excess EGF-TMR was retained in the sample during imaging enabling observations over incubation times from 3-60min. We observed a mean of ~57 EGFR tracks per cell across all incubation times from 117 cells, similar to the ~66 tracks per cell observed when EGF was absent, from 19 cells (table S1). Colocalization of EGFR and EGF foci was determined using numerical integration between overlapping green/red foci(27).

After EGF incubation from a few minutes, colocalization between green/red foci was detected (fig. 3A, movie S2 and fig. S5A). We estimated a mean ~15 foci per cell (40±18% of foci) were colocalized EGF-EGFR for 3-60min incubation. This value corresponds to 64% of all tracked EGFR molecules (fig. 3B,C). Colocalized EGF-EGFR foci had a higher mean stoichiometry (Student’s *t-*test P<0.0001) of 31 (table S1) compared to unligated clusters whose mean was 11, consistent with measurements made before adding EGF (fig.3D). The mean stoichiometry of unligated EGFR clusters remained roughly constant at 8-14 during incubation with EGF (fig.3E), compared to that of colocalized EGF-EGFR of approximately 20-40. Total EGFR-GFP copy number on the cell surface remained broadly constant after EGF was added (fig. S5C), implying that we mostly measure steady-state of endocytosis and recycling processes(29). Results from our kinetics model, discussed below, support this.

**Figure 3.**
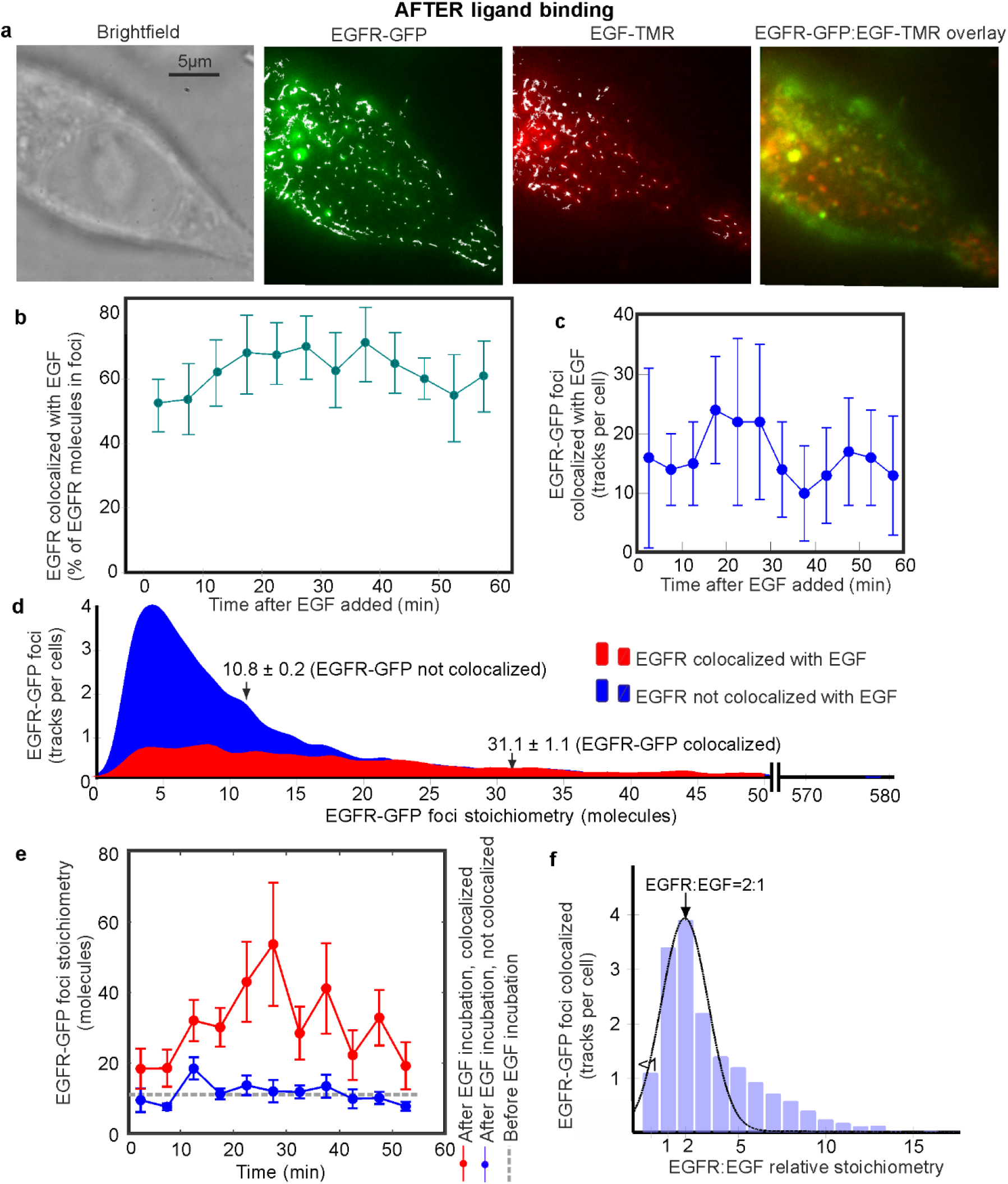
EGF effect on EGFR stoichiometry. **(A)** Brightfield and TIRF of SW620-EGFR-GFP after adding EGF (~10min time point), GFP (green), TMR (red) and overlay images shown (yellow indicates colocalization), tracking shown in white. **(B)** % of EGFR tracks colocalized to EGF, **(C)** number of colocalized EGF-EGFR tracks detected per cell (s.d. error bars). **(D)** Colocalized EGF-EGFR stoichiometry (red) and isolated foci (blue) across all times, mean and s.e.m. indicated, and **(E)** *vs.* time (s.d. error bars). Cells categorized into 6min interval bins, N=6-12 each bin. **(F)** Distribution of relative EGFR:EGF stoichiometry, overlaid Gaussian indicated, N=119 cells.

EGF-TMR *in vitro* using conjugation to coverslips exhibited step-wise photobleaching similar to GFP (fig. S3B). To determine the relative stoichiometry between EGFR and EGF when EGF was bound we measured red foci stoichiometry simultaneously to colocalized green foci, revealing a peak relative stoichiometry for EGFR:EGF of 1.9±0.8 (±half width half maximum, fig. 3F) with mean relative stoichiometry 4.2±0.1. The width of the fitted Gaussian underneath the 2:1 peak was consistent with variability in the GFP and TMR brightness we measured *in vitro*. Sub-dividing data by incubation time revealed no significant shift in relative stoichiometry from the 2:1 peak (fig. S5B). Before EGF-TMR was added in controls to the parental strain we detected rare autofluorescent foci resulting in pseudo colocalization of 2-3 tracks per cell (<3% of colocalized tracks), resulting in a small peak for apparent relative stoichiometry in green:red channels of ~0.5:1 (fig. S5D) thus having negligible impact on the 2:1 peak. Adding EGF-TMR to this strain resulted in the appearance of random foci in the red channel indistinguishable to that in the absence of EGF-TMR (Student’s *t-*test P>0.05).

Our observation of a peak EGF:EGFR stoichiometry ratio of 2:1 (mean 4:1) can be interpreted with a multi-state time-dependent kinetics model. Under the conditions of our experiments of relatively high EGF concentration, where we likely saturate EGFRs at the surface, the rate of internalization is 3-10%/min, dependent on cell line, and lower than at lower EGF concentrations owing to clathrin endocytosis pathway saturation. Recycling rates of ligand-occupied EGFR are ~10%/min, with recycling contributing significantly to the overall receptor distribution only after a pool of endosomal EGFR is accumulated.

Our time-dependent model shows that on adding EGF, initial concentrations of unligated EGFR monomers ([R]) and dimers ([RR]) decrease while concentrations of ligated monomers ([RL]) and dimers (singly ligated [RRL] and doubly ligated [RRL2]) increase over the first 5min (fig. 4a). Endocytosis leads to accumulation of internalized ligated monomers ([RL^inside^]) and dimers ([RRL^inside^] and doubly ligated dimers [RRL2^inside^]) (dashed lines, Fig.4A) with EGFR recycling back to the plasma membrane contributing to equilibration of all concentrations after 30-40min (fig. 4A). The fractional saturation on the surface Y^surface^, (ratio of EGF:EGFR on the surface, excluding internalization) is inset on fig. 4A, its inverse at equilibrium predicting EGFR:EGF in the absence of cooperativity of ~1.5, significantly lower than our mean ~4 (~2 peak value). However, if we correct for the temperature of our experiments (37°C) and assume negative cooperativity, as previously reported(22), our model predicts Y^surface^ ~0.24 which agrees with our experimental mean (i.e.~1/4) (fig. 4B). In light of our predictions, we can account for our data by a combination of negative cooperativity of binding, decreased affinity of ligand for dimers and reduced homo-dimerization on-rates (supplementary methods). These predictions could be consistent with initial EGF binding to monomeric EGFR to generate an activated state predisposed to dimerize with unligated EGFR. Our model, which accounts for recycling and endocytosis, enables rich interpretation of imaging data revealing insights that could not be achieved with time-independent models based solely on affinities and equilibrium constants. Fig. 4C shows the contrast between EGF:EGFR versus ligand concentration predictions at 37°C and 4°C; this arises from the strong temperature dependence of receptor internalization and of receptor ligand binding and dimerization equilibrium constants (full details see supplementary methods).

**Figure 4.**
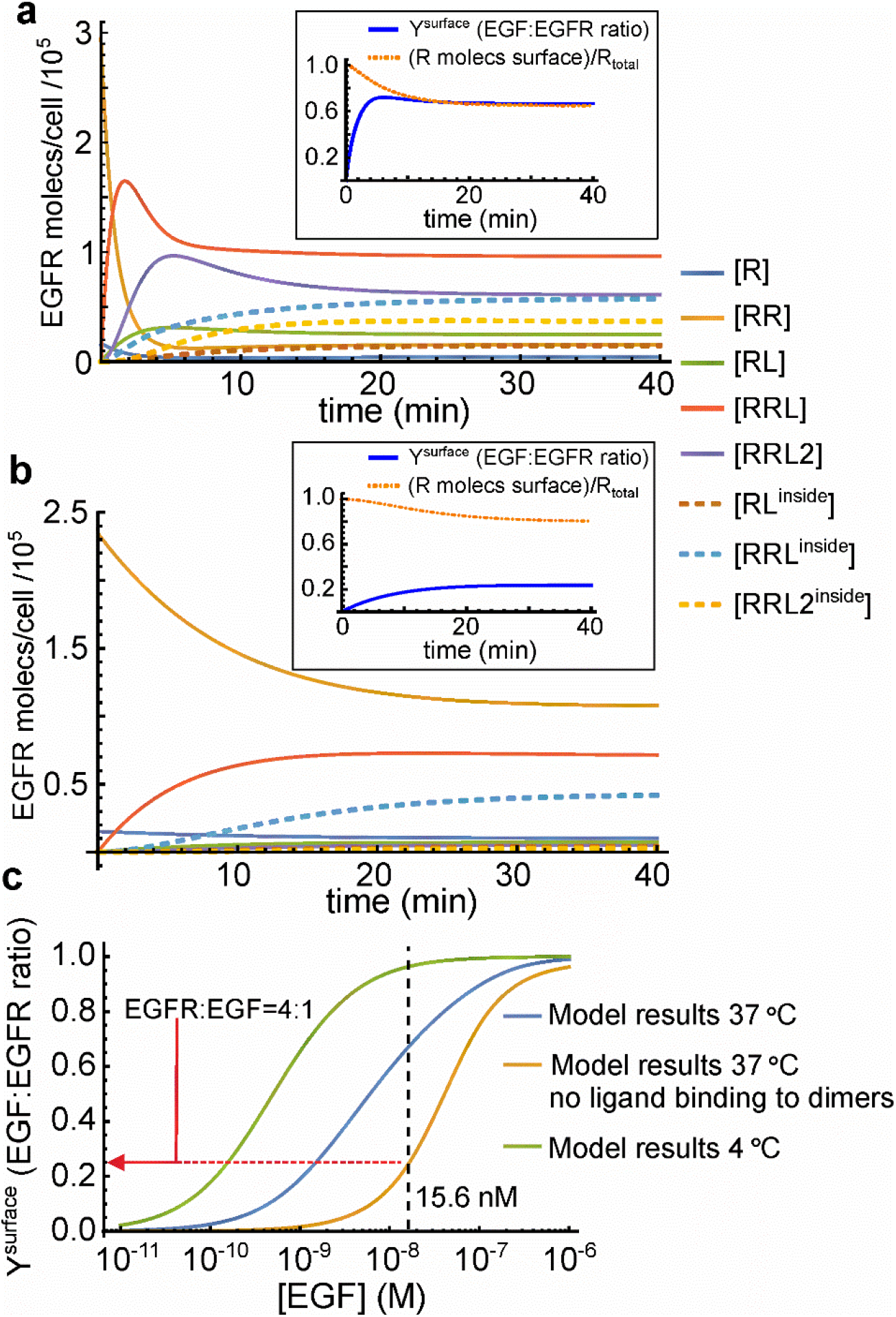
Time-dependent EGFR-EGF binding model. **(A)** Results of kinetics model for ligand binding to receptor monomers/dimers, dimerization, and endocytosis and recycling. Time dependence of receptor concentrations choosing parameters corresponding to 37°C. Inset: Y^surface^. **(B)** Predictions for same parameters of (A) but assuming ligand binds only to monomers. **(C)** Equilibrium Y^surface^ versus EGF concentration for parameters shown in (A) and (B), and those at 4°C. Black dashed line: experimental EGF concentration, red line: equivalent value of EGFR:EGF that we measure experimentally.

### EGFR clustering affected by EGF inhibition

To further understand the effect of EGF binding on EGFR clustering we performed TIRF in the presence of cetuximab or trastuzumab. Cetuximab is a monoclonal antibody anti-cancer drug commonly used against neck and colorectal cancers in advanced stages to inhibit cell division and growth(30). It binds to domain III of the soluble extracellular region of EGFR which is believed to result in partial blockage of the EGF binding region. This binding is believed to inhibit the receptor from adopting an extended conformation required for EGFR dimerization. Trastuzumab is a monoclonal antibody anti-cancer drug, commonly used to treat breast cancer(31); it has similar effects of inhibiting cell division and growth, however, it does not bind directly to EGFR but to domain IV of the extracellular segment of HER2(32) that does not affect HER2 self-association(33) but influences the stability of HER2-mediated dimers with EGFR(34).

Before adding EGF we found that treatment with cetuximab or trastuzumab at cytostatic concentrations resulted in significant increases in the mean EGFR-GFP stoichiometry of 25% and 65% (Student’s *t-*test, P<0.0001) respectively (fig. 5A), but with no significant effect on the number of detected EGFR-GFP tracks per cell. Adding EGF resulted in ~20% fewer colocalized EGF-EGFR tracks for cetuximab-or trastuzumab-treated cells compared to untreated cells (fig. 5B).

**Figure 5.**
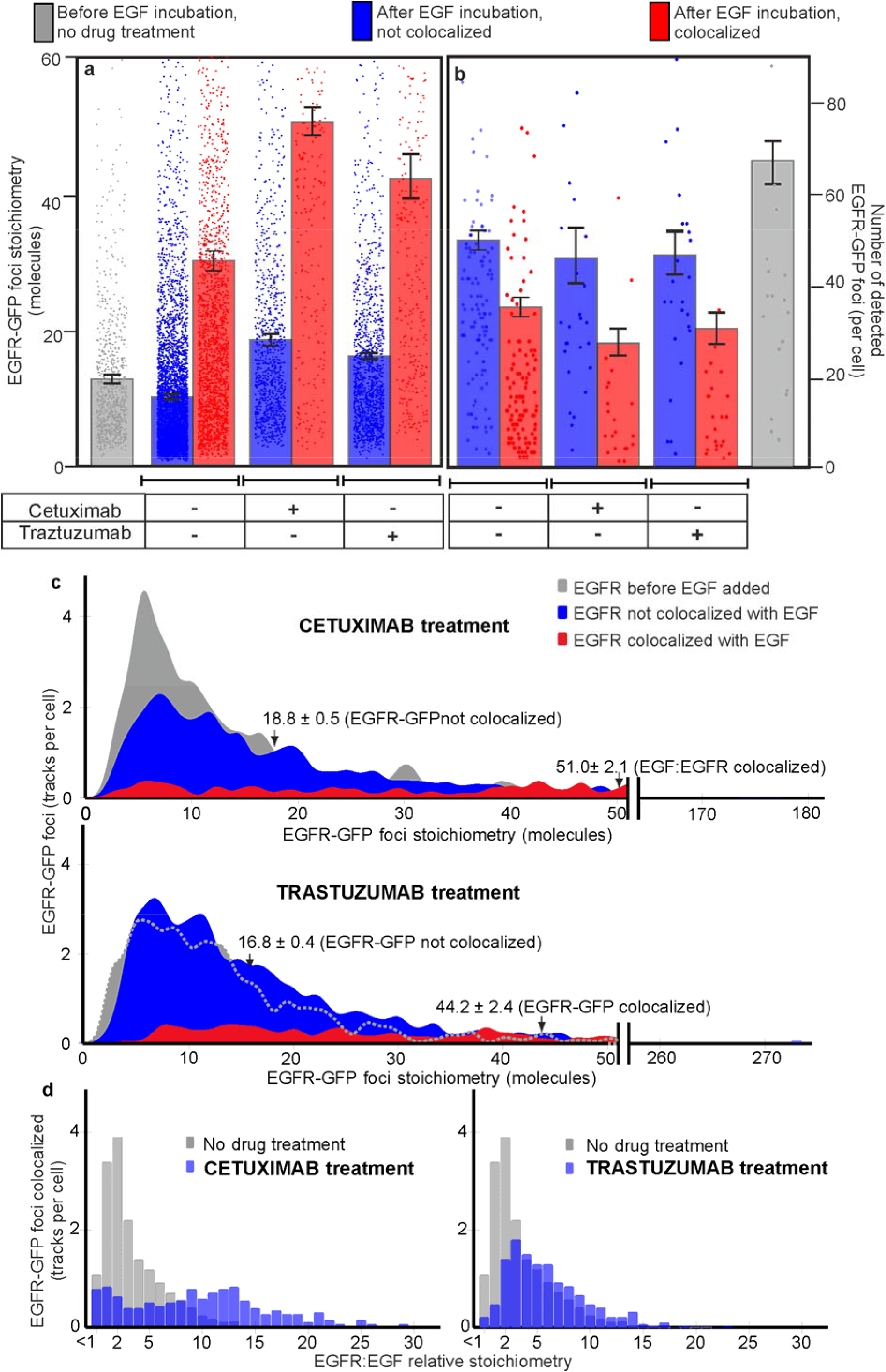
Effect of cetuximab and trastuzumab on EGF colocalization with EGFR. **(A)** Variation of mean EGFR-GFP foci stoichiometry, and **(B)** number of EGFR-GFP foci detected per cell. Colocalized EGF-EGFR (red) and isolated EGFR foci (blue) indicated for ± addition of cetuximab and trastuzumab. Error bars s.d, N =10-117 cells per dataset. **(C)** Distribution of EGFR foci stoichiometry for cells treated with cetuximab or trastuzumab, showing pre (grey) and post EGF addition for colocalized EGF-EGFR (red) and isolated EGFR (blue) foci, data collated across 60min EGF incubation, mean and s.e.m. indicated. **(D)** EGFR:EGF relative stoichiometry of colocalized EGF-EGFR foci for drug-treated cells (blue) contrasted against no drug treatment (gray). N=10-117 cells per dataset.

The mean colocalized EGF-EGFR foci stoichiometry in cetuximab and trastuzumab treatments was 51±2 and 44±2 respectively, with maxima of several hundred (fig. 5A,C). We also observe a shift to higher EGFR:EGF relative stoichiometry for cetuximab and trastuzumab treatments beyond the 2:1 peak observed for untreated cells (fig. 5D). The collapse of the peak at lower EGFR:EGF under cetuximab treatment reflects competitive binding with EGF. We also tested the inhibitor pertuzumab, a similar drug to trastuzumab albeit with complementary function against HER2/HER3 heteroassociation(35). Stoichiometry distributions were similar (fig. S9) to trastuzumab but full characterization is the subject of further study.

### EGF triggers larger EGFR heterocluster formation

Tracking of EGFR foci indicated Brownian diffusion up to time intervals of approximately 100ms (fig. 6A), while at longer times (fig. S6A) exhibiting transiently confined diffusion into zones of diameter 400-500nm (time intervals 100-600ms), and Brownian diffusion (time intervals >600ms). Using the initial gradient of the mean square displacement with respect to time interval for each track we determined the diffusion coefficient *D* and correlated this against EGFR foci stoichiometry. We used a simple model based on the Stokes-Einstein relation, in which the cross-sectional area of an EGFR cluster parallel to the plasma membrane scales linearly with the number of EGFR dimers present. The model assumes that *D*=*k_B_T*/*γ* where *k_B_* is Boltzmann’s constant, *T* absolute temperature and *γ* drag of the cluster in the membrane. Drag is proportional to the effective radius of the cluster, implying *D* is proportional to the reciprocal of the square root of the stoichiometry. Our model results in reasonable agreement for data corresponding to pre and post EGF incubation (fig. 6B).

**Figure 6.**
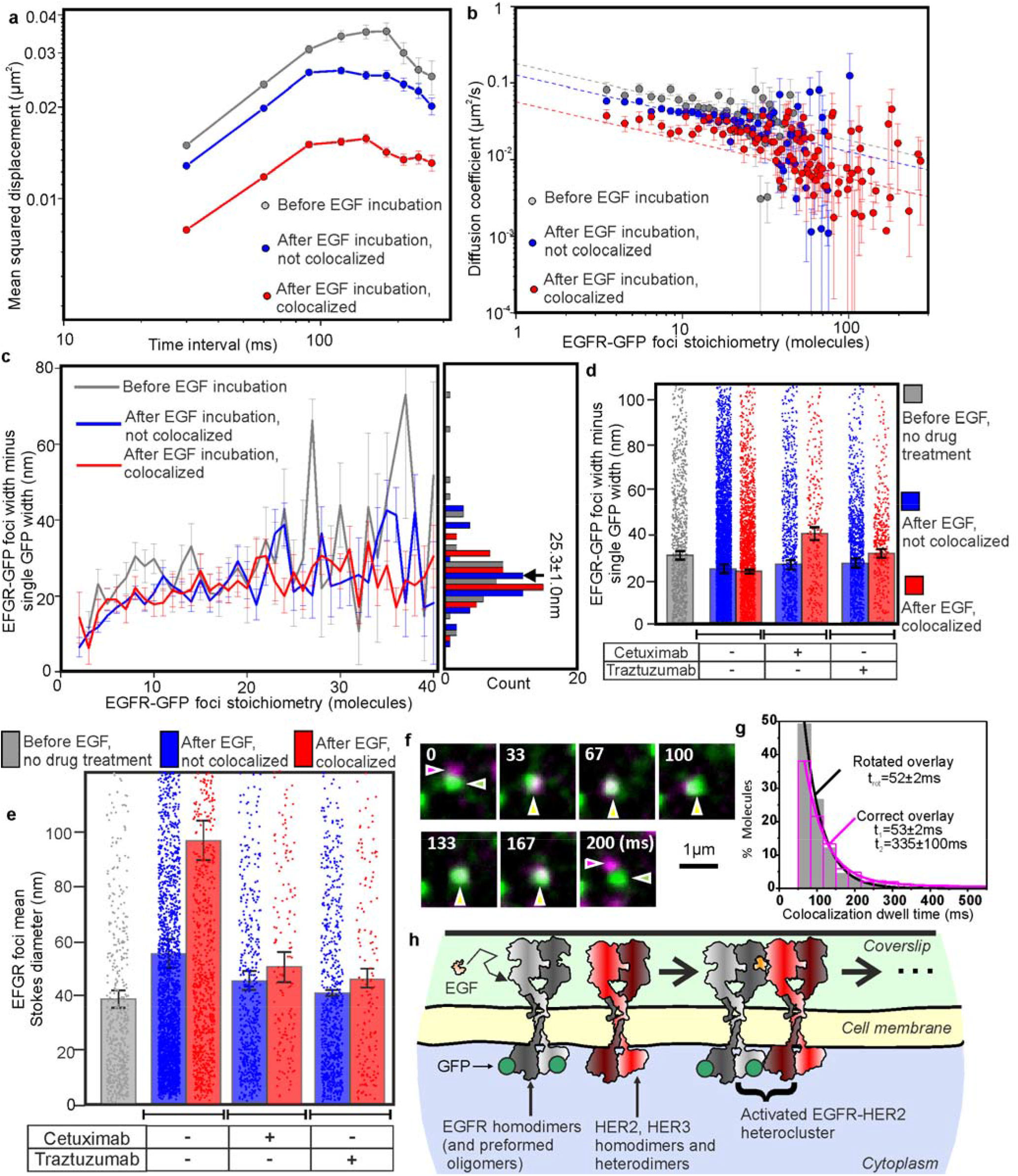
EGFR mobility depends on stoichiometry and EGF binding. **(A)** Log-log plot for average mean squared displacement for time intervals ≤300ms;**(B)** log-log plot for diffusion coefficient *D* with EGFR stoichiometry *S*, fits to Stokes-Einstein model *D*~*S*^−1/2^ (dashed lines). **(C)** EGFR-GFP foci minus single GFP width *vs.* stoichiometry, mean for all datasets indicated. Pre-EGF incubation (gray, N=770 foci, N=19 cells) and post EGF incubation for colocalized EGF-EGFR (red, N=1,969 foci, N=117 cells) and isolated EGFR (blue, N=1,741 foci, N=117 cells) shown. **(D)** Histograms of EGFR-GFP foci minus single GFP width. Pre EGF incubation for cells untreated with drugs (gray, N=1,252 foci, N=19 cells); cetuximab-treated cells post EGF incubation for colocalized EGF-EGFR (red, N=151 foci, N=10 cells) and isolated EGFR (blue, N=1,253 foci, N=10 cells) shown; trastuzumab-treated cells post EGF incubation for colocalized EGF-EGFR (red, N=263 foci, N=27 cells) and isolated EGFR (blue, N=1,479 foci, N=27 cells) shown. Errors s.e.m. **(E)** Histogram of mean Stokes diameter upon drug treatment, same datasets as for fig. 6D, s.e.m. error bars. **(F)** Single-molecule TIRF of EGFR-HaloTag650 (magenta arrows) and HER2-GFP (green arrows) undergoing transient colocalization and co-diffusion (yellow arrows), time since start indicated. **(G)** Histogram for the dwell time of colocalized EGFR–HER2 foci. The random colocalization dwell times were estimated by measuring the apparent colocalization between the green and red channels, after the red channel was rotated by 180°. The histogram for these was fitted well by a single exponential with time constant t_rot_ (magenta), whereas the colocalization dwell time histogram for which there was no prior rotation of the red channel (gray) was significantly different as (Brunner-Munzel test P<0.05) and required the sum of two exponentials for a reasonable fit with time constants t_1_ and t_2_. The t_1_ parameter was within error of t_rot_, which we assign as random colocalization, while t_2_ was assigned to non-random colocalization of EGFR and HER2. 285 random and 400 non-random colocalization events detected, N=4 cells. **(H)** Schematic illustrating how HER2/HER3 and EGFR dimers might associate following EGF ligation

We quantified EGFR-GFP foci widths by performing intensity profile analysis on background-corrected pixel values(26) and compared this with measurements from single GFP *in vitro*, as a function of stoichiometry *S* (fig. 6C). The mean EGFR-GFP foci width was greater than that of single GFP, which increased with *S*, consistent with a spatially extended structure. The dependence of this increase could be modelled with a heuristic power law *S^a^* with exponent *a*=0.27±0.04 (s.e.m.) showing no dependence with EGF ligation (fig. S6B), with mean EGFR-GFP foci minus single GFP width for all data of 25.3±1.0nm (s.e.m.). At the low end of *S* the increase in foci minus single GFP width was ~11-12nm, while at the high end, corresponding in some cases to several hundred EGFR, the increase in width was 30-40nm. Foci widths indicated no significant differences upon addition of cetuximab or trastuzumab prior to addition of EGF (P>0.05), however, we observed an increase of ~50% for EGF-EGFR foci for cetuximab-treated cells (P<0.001) (fig. 7D). Cells treated with cetuximab or trastuzumab exhibited a similar shape for mean square displacement *vs*. time interval to untreated cells (fig. S7A). Both treatment groups showed reasonable agreement to a Stokes-Einstein model, with/without EGF (fig. S7b).

We used *D* to estimate the physical diameter of EGFR foci. A full analytical treatment models diffusion of membrane protein complexes as cylinders with their long axis perpendicular to the membrane surface requiring precise knowledge of local membrane thickness, however, here we simplified analysis by calculating the diameter of the equivalent Stokes sphere to generate indicative values of drag length scale. We approximated drag as 3π*ηd* where *d* is the sphere diameter, assuming contributions from extracellular and cytoplasmic components are negligible since the kinematic plasma membrane viscosity *η* is higher by 2-3 orders of magnitude. Using a value of ~270cP estimated from human cell lines using high precision nanoscale viscosity probes(36), indicates a mean diameter of 40-60nm for isolated EGFR. Colocalized EGF-EGFR foci had a mean diameter closer to ~90nm, reduced back to the isolated EGFR levels within experimental error upon treatment of cetuximab or trastuzumab (fig. 6E).

The Stokes diameter for EGFR clusters is a measure of fluorescent EGFR-GFP plus any unlabeled components contributing to drag. Here, the endogenous level of unlabeled EGFR is low. However, other studies suggest that EGFR forms heterocomplexes with other RTKs as well as recent evidence of a HER2 inhibitor lapatinib inducing HER2/HER3 heterocomplex formation in breast cancer cells(37), although the expression of HER2, HER3 and HER4, is also low. However, inclusion of HER2 in these complexes was evidenced further by performing TIRF on CHO-K1 cells with similar low endogenous EGFR expression. We constructed a dual-label cell line containing GFP labelled HER2 and EGFR labelled with HaloTag650 (HaloTag STELLA Fluor 650) ligand (Methods). Using similar TIRF we found that HER2 and EGFR exhibit mobile and immobile foci, with mobility enabling transient colocalization and co-diffusion (fig. 6F over a mean non-random dwell time of 335±100ms (fig.6G, fig. S8A-C, movies S3,S4). The distributions of foci brightness for EGFR and HER2 were significantly greater than those measured for a single dye *in vitro*, consistent with a range of cluster stoichiometries beyond purely monomeric (fig. S8D).

Since the mean diameter of EGF-EGFR foci of ~90nm corresponds to a stoichiometry of approximately 16 EGFR dimers, the average diameter associated with a single dimer which accounts for the same cluster area is ~20nm, greater than the measured diameter of an EGFR dimer from crystal structures by a factor of 2. In other words, the observed diameter might be explained if EGFR-GFP dimers associate in a 1:1 relative stoichiometry with unlabeled dimers, presumably HER2 or HER2 associated with HER3, of similar size and structure. Although expression levels of these potential EGFR partners are low (fig. S1), only a small proportion of total cell EGFR is observed in clusters and colocalized with EGF (table S1). We observed 17 colocalized foci per cell containing a mean of 527 EGFR-GFP molecules in total (table S1). However, since TIRF only excites ~1/3 of the whole cell surface this indicates that there are approximately 1,500 EGFR-GFP molecules colocalized with EGF in total, ~0.7% of the cell copy number. A proportion of 0.7% of the expressed mRNA for EGFR-GFP following addition of EGF is at level comparable to the expressed mRNA for HER2 (fig. S1). The expressed mRNA corresponding to unlabeled EGFR is also at similar levels but we have no evidence that this is incorporated preferentially in EGFR-GFP:EGF clusters. Therefore we believe heterodimers with HER2, or HER2 associated with HER3, are the most likely explanation.

An additional phenomenon to consider is plasma membrane invagination as EGFR clusters grow, culminating in clathrin-coated cytoplasmic vesicles. Since visible foci detected in TIRF correspond to GFP localization in the invaginated basal membrane projected laterally onto our detector, their visible diameter might appear to approach an asymptotic plateau with respect to EGFR-GFP stoichiometry, broadly what we observed (fig. 6C).

## Discussion

Our findings from genetics, cell biology, biochemistry and biophysics, in particular single-molecule TIRF with super-resolved tracking, on live bowel carcinoma cells, suggest preformed homo-oligomeric EGFR is present in the plasma membrane prior to EGF ligation, comprising predominantly clusters of EGFR dimers (fig. 6B). We chose a bowel carcinoma cell line which does not natively express EGFR, rather than use CRISPR/Cas9 to modify a natively EGFR expressing carcinoma cell line which may have also co-evolved different expression patterns, complicating our observations of EGFR behavior. Using GFP on EGFR with TMR on EGF enabled insight into stoichiometry, mobility and kinetics of single EGFR clusters in their pre and post ligation states. Our observations indicate the most prevalent tracked EGFR oligomer in the absence of bound EGF is a hexamer, though with higher order oligomers present extending to ~90 molecules. We find that EGF ligation results in higher stoichiometry, contrary to earlier reports suggesting tetrameric EGFR is the most likely state(19). We observe that commonly used anti-cancer drugs result in changes to the EGFR content of clusters. By comparing the mobility of ligated EGFR clusters we measured cluster diameters, indicating that EGF ligation results in formation of heteroclusters containing a mixture of EGFR and HER2, or HER2 associated with HER3. These observations were consistent with TIRF on transfected CHO-K1 showing EGFR and HER2 transiently interacting over several hundred milliseconds even before EGF ligation. Using a multi-state kinetics model which investigates time-dependent EGF-EGFR interactions we find our observations are consistent with predictions based on negative cooperativity, preferential binding of EGF ligand to EGFR monomers and preferential dimerization of ligated-unligated monomers. Two important improvements in our study over earlier reports are that: (i) our findings relate to a primary human carcinoma cell strain, enabling insights to the EGF pathway in cancer directly; (ii) we have definitive spatial information concerning EGFR and EGF localization simultaneously and so have confidence concerning the effects of EGF ligation on the stoichiometry of specific EGFR foci. In prior microscopy in which labelled EGF is not imaged simultaneously to labelled EGFR inference is more limited.

Our findings show EGFR is clustered before and after EGF ligation, consistent with observations from earlier AFM using EGF-coated tips which probed the surface of human lung adenocarcinoma cell line A549, known to have high EGFR expression(38). This study suggested half the EGFR clusters had pre-activated diameters 20-70nm, 35-105nm post activation, comparable with our measurements. However, we find important differences with respect to some previous single-molecule studies. Although there were earlier suggestions of preformed EGFR oligomers, Needham et al(19) and Huang et al(20), report putative monomeric EGFR, in particular Huang et al assign a high apparent monomeric proportion of 94%. We cannot directly exclude that monomeric EGRF are present at such high surface density in our experiments that their mean separation is less than the optical resolution, thus untrackable. However, the absence of not a single detected monomer from several thousand tracks from all datasets, despite having the sensitivity to detect single GFP (fig. S4C), makes this explanation unlikely. A more plausible explanation may lie in differences in copy number; in our experiments we estimate ~200,000 EGFR molecules per cell similar to endogenously expressing cancer cell lines(39) but more than double that estimated from Needham et al and Huang et al, which may account for shifting the equilibrium position for EGFR oligomerization towards higher stoichiometries. This upshift in oligomer formation on-rate may also contribute to a depleted monomeric EGFR population in our observations, which has implications for several carcinomas in which the expression level of EGFR is known to be high.

Our peak value of 6 EGFR before EGF ligation cannot be explained by a model as proposed by Needham et al suggesting face-to-face dimers associate with the EGFR dimer interface between back-to-back dimers to generate higher order complexes; their model predicts a most likely stoichiometry of 4, and EGFR oligomers as extended structures which would in principle manifest as *D*~*S*^−1^, whereas our mobility analysis suggests a dependence of *D*~*S*^−1/2^. As discussed above, differences in copy numbers may partially explain a shift in stoichiometry to higher values. The physical driving force behind cluster formation is something we do not directly address here, however, there is evidence that forces associated with molecular crowding in the membrane may result in oligomerization of proteins and the appearance of complex cytoskeletal and clathrin pit morphologies, as well as electrostatic protein-lipid (40) and direct protein-protein interactions(41) being possible contributory factors towards EGFR cluster generation.

Earlier work on heterocomplex formation showed EGFR may associate with other ERBB proteins including HER2(16), however, there are discrepancies as to whether these associations are before or after EGF ligation. Our observations suggest heterocomplex formation increases following EGF ligation. Our findings that HER2-dimerization inhibitor trastuzumab influences the stoichiometry of ligated EGFR clusters might indicate a role for this drug in modulating regulatory balance through the availability of endogenous HER2 to associate with EGFR, though our experiments cannot directly exclude the presence of HER3 also. Even when scarce, the presence of HER2 is known to selectively discourage internalization and degradation of activated EGFR, and promote recycling to the plasma membrane both via chaperone proteins and EGF dissociation(42). The physiological role of heterocomplex formation is unclear. HER2 is known to act as coreceptor but has no known direct ligand. The mobility of heterocomplexes may enable a spread of signal across cell surfaces, especially if HER2 turns over between EGFR complexes as suggested by transient colocalization between HER2 and EGFR in CHO-K1. One consequence of HER2 association after EGF binding is that the whole cell signal response is more likely to be highly biphasic. The resistance of HER2-bearing complexes to downregulation also acts to sustain signaling once established. Our findings of increases in heterocomplex cluster size post EGF ligation may suggest new strategies for anti-cancer drug design. For example, new drugs to target interaction interfaces between HER2 and EGFR directly. Alternatively, it may be valuable to explore new strategies to disrupt the oligomeric nature of EGFR before EGF ligation. Similarly, there may be value in using our single-molecule quantification to investigate different human carcinomas, for example those of the lung in which EGFR mutations are implicated in cancer(43). With these future studies there may also be value in pursuing CRISPR based gene-editing technologies for generating fluorescent fusions to mitigate against the risks of increased levels of unlabeled endogenous EGFR using conventional transfection methods which retain the native gene. Also, in enabling robust quantification of the actions of different cancer drugs there may be value in enabling future insights as to relative doses of each that are most efficacious in chemotherapy (i.e. a dose ‘sweet-spot’) in carcinomas known to be treatable using combined drugs, such as in gastric cancer(44).

## Materials and Methods

### Cell lines

Colorectal carcinoma line SW620 and CHO-K1 were both stably transfected with fluorescently tagged EGFR and HER2 using standard methods. Full details in supplementary methods.

### RT-qPCR

To extract RNA, cell pellets were lysed in Trizol (Invitrogen). RNA was converted into cDNA using MMLV reverse transcriptase (New England Biolabs®) with Oligo(dT)12-18 primers (Invitrogen), 10mM dNTP mix and RNase inhibitor Ribolock (Thermo Fisher Scientific), cDNA purified using QIAquick PCR purification (QIAGEN). Expression levels of *HER2*, *HER3*, *HER4* and *EGFR* were determined by qPCR using Fast SYBR Green Master Mix on QuantStudio TM 3 Real-Time PCR System (Thermo Fisher Scientific), 20s/95°C then 40 cycles of 1s/95°C and 20s/60°C, normalized against housekeeping *PLQC2*. Relative fold expression change was calculated using ΔΔCt analysis.

### Microarray

Gene expression data for 78 unique, non-duplicate (not sourced from same patients) colorectal cancer cell lines were obtained by performing microarray using the Affymetrix GeneChip HG-U133 Plus 2.0 microarray, normalized using RMA and batch-removed using Partek Genomics Suite software. Full details in supplementary methods.

### Fab

IgG antibodies to EGF and anti-EGF rabbit anti-mouse polyclonal IgG (Molecular Probes) were digested by papain, confirmed by migration of 28-30kDa and 25kDa proteins corresponding to reduced Fc and Fab respectively. Fab was purified using protein A immobilized within a spin column, evaluated by 280nm absorbance (Thermo Scientific NanoDrop).

### Confocal

Zeiss inverted Axio Observer Z1 microscope with LSM 510 META scanning module and Plan-Aprochromat 63x 1.40NA oil immersion DIC M27 objective lens was used, enabling simultaneous imaging of green/red channels via 488nm/565nm wavelengths. SW620:EGFR-GFP cells grown in Corning 75cm^2^ treated plastic cell culture flasks in a humidified incubator (37 ºC, 5% CO_2_) once 70-100% confluent were subcultured by trypsinization. 2-7 days prior to imaging, ~200,000 cells were seeded onto a Ibidi μ-dish 35mm, high glass bottom using their normal culture media, DMEM, containing phenol red, then changed to DMEM with addition of 4.5g/l glucose, L-glutamine, HEPES, without phenol red, and supplemented with 10% FBS, 100 units/ml penicillin and 100μg/ml streptomycin, or directly into DMEM without phenol red as appropriate. Prior to imaging media was changed to Molecular Probes® Live Cell Imaging Solution supplemented with 1.5mg/ml G418 sulfate.

For immunofluorescence we harvested SW620-EGFR-GFP cells 48h prior to fixation at ~50,000 density per well seeded into Ibidi μ-Slide VI0.4, cultured in DMEM without phenol red, supplemented with 4.5g/l glucose, L-glutamine, HEPES, 10% FBS and 100 units/ml of penicillin and 100μg/ml streptomycin, 1.5mg/ml G418. Cells were fixed with 4% formaldehyde at room temperature for 10min and washed. Non-specific antibody adsorption was blocked with 10% FBS in PBS for 10-20min. Primary antibodies were EGFR (D38B1) XP rabbit monoclonal 4267P (Cell Signaling Technology, 1:50 dilution) and anti-GFP chicken IgY (H+L) (Cell Signaling Technology, 1:400 dilution) in PBS with 10% FBS and 0.1% saponin overnight at 4 ºC. Each well was washed with 10% FBS and incubated with secondary antibodies, DyLight 633 goat anti-rabbit immunoglobulin G (IgG) highly cross adsorbed (PN35563, Thermo Scientific), 1:200, and Alexa Fluor 633 goat anti-chicken IgG (H+L) 2 mg/ml (Invitrogen) in PBS with 10% FBS and 0.1% saponin. Channels were washed with PBS and Sigma Aldrich Mowiol 4-88 added to solidify overnight. GFP, DyLight 633 or Alexa Fluor 633 and 4’,6-diamidino-2-phenylindole (DAPI) were individually illuminated and scanned (indicating no mycoplasma). GFP was excited as for live cell imaging, while DyLight 633 and Alexa Fluor 633 were excited by a 633nm HeNe laser.

### TIRF

For SW620:EGFR-GFP a dual-color single-molecule microscope was modified from a previous design(27) equipped with nanostage (Mad City Labs) and 37 ºC humidified incubator supplemented with 5% CO_2_ (INUB-LPS, Tokai Hit). We used Elforlight B4-40 473nm 40mW and Oxxius SLIM 561nm 200mW lasers attenuated into a common path prior to polarization circularization (achromatic λ/4 plate) before entering a Nikon Eclipse-Ti inverted microscope body. An achromatic lens mounted onto a translation stage controlled the angle of incidence into the objective lens to generate TIRF via a Semrock 488/561nm BrightLine® dual-edge laser-flat dichroic beam splitter into a Nikon TIRF 100x NA1.49 oil immersion objective lens enabling simultaneous GFP/TMR detection across a 20μm full width at half maximum field, intensity 1kW/cm^2^, 100nm penetration depth. Fluorescence was sampled 30ms per frame imaging onto two 512×512 pixel array EMCCD cameras (Andor, iXon+ DU-897 and iXon DU-887 for green/red, piezoelectrically cooled to −70ºC), 50nm/pixel magnification, via Semrock 561nm StopLine® single notch and Chroma 473nm notch filters. Typically, scans were 200 frames. For *in vitro* TIRF we used surface-immobilized GFP or EGF-TMR via anti-GFP or anti-EGF antibodies (Molecular Probes) or Fab followed by BSA passivation prior to washing(24). Slides were constructed from Ibidi sticky-Slides VI0.4 and 25mm×75mm No. 1.5 D263M Schott plasma-cleaned glass coverslip and IgG/Fab applied to a single channel and incubated at room temperature for 5min, washed x3 PBS, blocked with 1mg/ml of BSA for 60min. The channel was again washed x3 then incubated with GFP for 7.5min or EGF-TMR for 4min. The channel was washed x5 before adding 1:10000, 200nm diameter, 4% w/v, Invitrogen Molecular Probes carboxyl latex beads for focusing.

For live cell TIRF, cells were seeded/grown in media onto glass-bottomed Petri dishes or Corning culture flasks at 37 ºC, 5% CO_2_. SW620:EGFR-GFP, or SW620 as negative control, imaged on either i) plasma cleaned glass coverslips (25mm×75mm No. 1.5 D263M Schott) covered by a sterile Ibidi sticky-Slide VI0.4, or ii) Ibidi μ-dish 35mm, high glass bottom as for confocal. 48h prior to imaging, cells were seeded onto the imaging chamber at ~200,000/cm^2^ density. For slides, 50μl (or 800μl for dishes) DMEM without phenol red supplemented with 10% FBS, 100 units/ml penicillin and 100μg/ml streptomycin was added. 24h prior to imaging media was changed to DMEM without phenol red supplemented with 100 units/ml penicillin, 100μg/ml streptomycin and 1.5mg/ml G418 sulfate plus inhibitors (2ng/ml cetuximab (BioVision), 10ng/ml trastuzumab (BioVision) or 20ng/ml pertuzumab (Selleck Chemicals)) where necessary, without FBS (starving cells of residual serum EGF) for 24h. We checked SW620 for expression of the most common ligands, using publicly available RNA-Seq data and our microarray data: EGF zero; TGFA low level; HBEGF low level expression; AREG zero; BTC zero; EREG zero; EPGN no data available. Although we cannot rule out the presence of very low levels of TGFA, cells were washed prior to imaging and no change was observed in EGFR clustering over 60min unless EGF was added (fig. 3) suggesting no or negligible autocrine EGFR stimulation. Immediately before imaging, media was exchanged to Molecular Probes® Live Cell Imaging Solution supplemented with G418 sulfate and inhibitors where appropriate. Fluorescence sequences at 5min intervals up to 60min were acquired after adding 100ng/ml (15.6nM) EGF-TMR (Molecular Probes).This EGF concentration resulted in clear phosphorylation activity on western blots and is consistent with high physiological levels found in prostate and breast tissue. Full details in supplementary methods.

CHO-K1 cells were illuminated using a different TIRF microscope with similar capability. Objective lens based excitation was used with an evanescent field of 100nm, and 37°C stage temperature control, around an IX-83, Olympus inverted microscope with Olympus 100× NA1.49 oil immersion objective lens, laser powers 1.2mW and 5mW for 488nm and 642nm lasers. Dual color images were separated by dichroic mirrors (ZT405/488/561/640rpc-UF3, ZT561rpc-UF3 and ZT640rpc-UF3; Chroma), projected into green/red detection channels with emission filters of 500–550nm for HER2-mGFP (ET525/50m; Chroma) and 662.5–737.5nm (ET525/50m; Chroma) for EGFR labelled with HaloTag STELLA Fluor 650 ligand (a red fluorescent dye), then onto a two-stage microchannel plate intensifier (C9016-02MERLP24; Hamamatsu Photonics), lens-coupled to a high-speed scientific complementary metal oxide semiconductor sensor camera (C1440-22CU; Hamamatsu), 33ms per frame. For fluorescence labelling of Halo7-tagged proteins, cells were incubated with 30nM STELLA 650-conjugated HaloTag ligand (GORYO) in Ham’s F12 media (Invitrogen), 37°C 20min, washed x3, and media replaced by Ham’s F12 media with 2mM PIPES, pH7.0.

### Tracking

For SW620:EGFR-GFP MATLAB (MathWorks)(27) code was used to track foci in green/red channels to determine spatial localization and calculate integrated pixel intensities and diffusion coefficients. The centroid of each focus was determined using iterative Gaussian masking to sub-pixel precision of 40nm, brightness calculated as the summed intensity inside a 5-pixel-radius centroid-centered circle, after subtraction of local background, signal-to-noise ratio (SNR) defined as intensity divided by background standard deviation. For SNR >0.3 (optimum for high true and low false positive detection from simulations trained on *in vitro* data) a focus was accepted and fitted with a 2D radial Gaussian to determine its sigma width. Foci detected in consecutive images separated by ≤5 pixels and not different in brightness or width by more than a factor of two were linked into the same track. For CHO-K1 foci tracking used a similar algorithm.

### Stoichiometry

Stoichiometry per track was estimated in MATLAB using step-wise fluorophore photobleaching to determine GFP or TMR brightness(24) from live cells and corroborated *in vitro*. Live cell foci brightness followed exponential photobleaching. As each focus photobleaches it will emit the characteristic single GFP or TMR brightness value, *I_GFP_* or *I_TMR_*, detected as the peak of foci intensities over time. Estimates for *I_GFP_* and *I_TMR_* were verified by Fourier spectral analysis(24) yielding the same value within error. Initial intensity *I_0_* was estimated by interpolation of the first 3 points in each track, stoichiometries by dividing *I_0_* by the single-molecule fluorophore brightness, distributions rendered as kernel density estimations(24).

### EGFR-EGF time-dependent kinetics

We developed a multi-state time-dependent kinetics model for ligand binding to receptor monomers and dimers, incorporating homo-and hetero-dimerization of ligated and unligated receptors, internalization of ligated receptors via endocytosis and subsequent recycling of receptors to the plasma membrane that solves multiple rate equations to determine concentrations of ligated and unligated receptor monomers and dimers, and concentrations of internalized receptors, as a function of time (full details supplementary methods).

### Software access

All bespoke code in MATLAB is available from EGFRanalyser at https://sourceforge.net/projects/york-biophysics/.

### Data availability

We do not upload additional data analysis files since analyzed data are included in full in the main text and supplementary files. All raw imaging data are available from the authors.

### Statistical analysis

Two-tailed Student’s *t-*tests were performed for comparisons between pairs of datasets to test null hypothesis that data in each was sampled from the same statistical distribution assuming (n_1_+n_2_-2) degrees of freedom where n_1_ and n_2_ are the number of data points in each distribution and by convention that *t* statistic values which have a probability of confidence P>0.05 are statistically not significant. For TIRF each cell was defined as a biological replicate sampled from the cell population with sample sizes of 10-117 cells per condition. Technical replicates are not possible with irreversible photobleaching, nevertheless. Differences between colocalization dwell times were assessed using the Brunner-Munzel rank order test.

## Acknowledgements

We thank Philippe Bastiaens, Max Planck Institute of Molecular Physiology, Dortmund, Germany for donation of plasmid perbB1-EGFP-N1, Ivan R. Nabi, University of British Columbia, Canada for donation of human EGFR-YFP plasmid, and Hannah Walker and Norman Maitland for technical advice concerning cancer cell maintenance and western blotting and for use of resources at the Cancer Research Unit, University of York.

## Author contributions

DO, JW, SS, CF created and biologically characterized the cell line. OH, ILG built the microscope. CF, ILG, OH, AL, PZ collected the microscopy data. AW, CF wrote analysis software. AW, CF, AH, PZ, TCL analyzed the data. ILG performed modelling. WB and MCL designed the study. All authors wrote the manuscript.

## Supplementary Materials

Supplementary Materials comprise movies S1-S4 plus a single compiled PDF containing supplementary methods, table S1, titles/legends of movies S1-S4, supplementary references, and figures S1-S9:

Table S1. Mean EGFR foci stoichiometry.
Fig. S1. EGFR expression levels.
Fig. S2. Confocal and TIRF characterization.
Fig. S3. Characterization of unitary fluorophore brightness values.
Fig. S4. More examples of cells before addition of EGF ligand.
Fig. S5. Random foci overlap model.
Fig. S6. Characterizing EGFR and EGF foci stoichiometry after addition of EGF
Fig. S7. EGFR foci diffusion.
Fig. S8. Treatment effects on mobility Fig. S8. EGFR and Her2 colocalization
Fig. S9. Effect of pertuzumab on EGFR foci stoichiometry
Movie S1. Live transected SW620 cell single-color TIRF imaging.
Movie S2. Live transected SW620 cell dual-color TIRF imaging.
Movie S3. Live CHO-K1 cell dual-color TIRF imaging.
Movie S4. Live CHO-K1 cell dual-color TIRF imaging zoom-in.

## References

1. R. Roskoski, The ErbB/HER family of protein-tyrosine kinases and cancer. Pharmacol. Res. 79, 34–74 (2014).

2. R. N. Jorissen, et al., Epidermal growth factor receptor: mechanisms of activation and signalling. Exp. Cell Res. 284, 31–53 (2003).

3. I. Lax, et al., Functional analysis of the ligand binding site of EGF-receptor utilizing chimeric chicken/human receptor molecules. EMBO J. 8, 421–7 (1989).

4. M. R. Schneider, E. Wolf, The epidermal growth factor receptor ligands at a glance. J. Cell. Physiol. 218, 460–466 (2009).

5. S. Cohen, R. A. Fava, Internalization of functional epidermal growth factor:receptor/kinase complexes in A-431 cells. J. Biol. Chem. 260, 12351–8 (1985).

6. M. A. Lemmon, et al., Two EGF molecules contribute additively to stabilization of the EGFR dimer. EMBO J. 16, 281–94 (1997).

7. M. Odaka, D. Kohda, I. Lax, J. Schlessinger, F. Inagaki, Ligand-binding enhances the affinity of dimerization of the extracellular domain of the epidermal growth factor receptor. J. Biochem. 122, 116–21 (1997).

8. T. Domagala, et al., Stoichiometry, kinetic and binding analysis of the interaction between epidermal growth factor (EGF) and the extracellular domain of the EGF receptor. Growth Factors 18, 11–29 (2000).

9. K. M. Ferguson, et al., EGF activates its receptor by removing interactions that autoinhibit ectodomain dimerization. Mol. Cell 11, 507–17 (2003).

10. J. L. Macdonald-Obermann, L. J. Pike, The Intracellular Juxtamembrane Domain of the Epidermal Growth Factor (EGF) Receptor Is Responsible for the Allosteric Regulation of EGF Binding. J. Biol. Chem. 284, 13570–13576 (2009).

11. P. Liu, et al., A single ligand is sufficient to activate EGFR dimers. Proc. Natl. Acad. Sci. 109, 10861–10866 (2012).

12. M. A. Lemmon, Ligand-induced ErbB receptor dimerization. Exp. Cell Res. 315, 638–48 (2009).

13. Y. Sako, S. Minoghchi, T. Yanagida, Single-molecule imaging of EGFR signalling on the surface of living cells. Nat. Cell Biol. 2, 168–72 (2000).

14. M. Martin-Fernandez, D. T. Clarke, M. J. Tobin, S. V Jones, G. R. Jones, Preformed oligomeric epidermal growth factor receptors undergo an ectodomain structure change during signaling. Biophys. J. 82, 2415–27 (2002).

15. A. H. A. Clayton, et al., Ligand-induced dimer-tetramer transition during the activation of the cell surface epidermal growth factor receptor-A multidimensional microscopy analysis. J. Biol. Chem. 280, 30392–9 (2005).

16. R.-H. Tao, I. N. Maruyama, All EGF(ErbB) receptors have preformed homo- and heterodimeric structures in living cells. J. Cell Sci. 121, 3207–17 (2008).

17. P. Nagy, J. Claus, T. M. Jovin, D. J. Arndt-Jovin, Distribution of resting and ligand-bound ErbB1 and ErbB2 receptor tyrosine kinases in living cells using number and brightness analysis. Proc. Natl. Acad. Sci. U. S. A. 107, 16524–9 (2010).

18. Y. Park, et al., Single-Molecule Rotation for EGFR Conformational Dynamics in Live Cells. J. Am. Chem. Soc. 140, 15161–15165 (2018).

19. S. R. Needham, et al., EGFR oligomerization organizes kinase-active dimers into competent signalling platforms. Nat. Commun. 7, 13307 (2016).

20. Y. Huang, et al., Molecular basis for multimerization in the activation of the epidermal growth factor receptor. Elife 5, e14107 (2016).

21. V. Palmieri, et al., Mechanical and structural comparison between primary tumor and lymph node metastasis cells in colorectal cancer. Soft Matter 11, 5719–5726 (2015).

22. J. L. Macdonald, L. J. Pike, Heterogeneity in EGF-binding affinities arises from negative cooperativity in an aggregating system. Proc. Natl. Acad. Sci. 105, 112–117 (2008).

23. J. L. Wilding, S. McGowan, Y. Liu, W. F. Bodmer, Replication error deficient and proficient colorectal cancer gene expression differences caused by 3’UTR polyT sequence deletions. Proc. Natl. Acad. Sci. 107, 21058–21063 (2010).

24. M. C. Leake, et al., Stoichiometry and turnover in single, functioning membrane protein complexes. Nature 443, 355–358 (2006).

25. A. J. M. Wollman, M. C. Leake, Millisecond single-molecule localization microscopy combined with convolution analysis and automated image segmentation to determine protein concentrations in complexly structured, functional cells, one cell at a time. Faraday Discuss. 184, 401–24 (2015).

26. A. J. M. J. Wollman, et al., Transcription factor clusters regulate genes in eukaryotic cells. Elife 6, e27451 (2017).

27. I. Llorente-Garcia, et al., Single-molecule in vivo imaging of bacterial respiratory complexes indicates delocalized oxidative phosphorylation. Biochim. Biophys. Acta 1837, 811–24 (2014).

28. M. C. Leake, D. Wilson, B. Bullard, R. M. Simmons, M. R. Bubb, The elasticity of single kettin molecules using a two-bead laser-tweezers assay. FEBS Lett. 535(2003).

29. A. Sorkin, J. E. Duex, Quantitative analysis of endocytosis and turnover of epidermal growth factor (EGF) and EGF receptor. Curr. Protoc. Cell Biol. Chapter 15, Unit 15.14 (2010).

30. P. Kirkpatrick, J. Graham, M. Muhsin, Fresh from the pipeline: Cetuximab. Nat. Rev. Drug Discov. 3, 549–550 (2004).

31. K. P. Garnock-Jones, G. M. Keating, L. J. Scott, Trastuzumab. Drugs 70, 215–239 (2010).

32. H.-S. Cho, et al., Structure of the extracellular region of HER2 alone and in complex with the Herceptin Fab. Nature 421, 756–760 (2003).

33. H. Maadi, B. Nami, J. Tong, G. Li, Z. Wang, The effects of trastuzumab on HER2-mediated cell signaling in CHO cells expressing human HER2. BMC Cancer 18, 238 (2018).

34. T. S. Wehrman, et al., A system for quantifying dynamic protein interactions defines a role for Herceptin in modulating ErbB2 interactions. Proc. Natl. Acad. Sci. U. S. A. 103, 19063–19068 (2006).

35. J. Rockberg, J. M. Schwenk, M. Uhlén, Discovery of epitopes for targeting the human epidermal growth factor receptor 2 (HER2) with antibodies. Mol. Oncol. 3, 238–247 (2009).

36. I. López-Duarte, T. T. Vu, M. A. Izquierdo, J. A. Bull, M. K. Kuimova, A molecular rotor for measuring viscosity in plasma membranes of live cells. Chem. Commun. 50, 5282–5284 (2014).

37. J. Claus, et al., Inhibitor-induced HER2-HER3 heterodimerisation promotes proliferation through a novel dimer interface. Elife 7(2018).

38. W. Zhao, et al., Mapping the resting and stimulated EGFR in cell membranes with topography and recognition imaging. Anal. Methods 6, 7689–7694 (2014).

39. F. Zhang, et al., Quantification of Epidermal Growth Factor Receptor Expression Level and Binding Kinetics on Cell Surfaces by Surface Plasmon Resonance Imaging. Anal. Chem. 87, 9960–9965 (2015).

40. Y. Wang, et al., Regulation of EGFR nanocluster formation by ionic protein-lipid interaction. Cell Res. 24, 959–976 (2014).

41. I. Chung, et al., Spatial control of EGF receptor activation by reversible dimerization on living cells. Nature 464, 783–7 (2010).

42. Z. Wang, L. Zhang, T. K. Yeung, X. Chen, Endocytosis Deficiency of Epidermal Growth Factor (EGF) Receptor–ErbB2 Heterodimers in Response to EGF Stimulation. Mol. Biol. Cell 10, 1621–1636 (1999).

43. J. G. Paez, et al., EGFR mutations in lung, cancer: Correlation with clinical response to gefitinib therapy. Science (80-.). 304, 1497–1500 (2004).

44. K. Aoyagi, et al., Molecular targeting to treat gastric cancer. World J. Gastroenterol. 20, 13741–13755 (2014).

