## Supplementary methods for "EGF signaling in bowel carcinoma cells utilizes higher order architectures of EGFR and HER2"

Title

^2^ Current address: Biosciences Institute, Newcastle University
NE2 4HHO, United Kingdom.

^4^ Current address: Okinawa Institute of Science and Technology Graduate University, 119-1 Tancha, Onna-son, Kunigami-gun, Okinawa, Japan 904-0495.

^5^ Current address: Department of Physics and Astronomy, University College London, Gower Street, London WC1E 6BT, United Kingdom

^6^ Current address: Department of Microbiology and Immunology, Institute for Biomedicine, Sahlgrenska Academy, University of Gothenburg, 405 30 Gothenburg, Sweden.

^7^ Membrane Cooperativity Unit, OIST, Onna-son, Japan.

^8^ MRC Weatherall Institute of Molecular Medicine, University of Oxford, John Radcliffe Hospital, Oxford OX3 9DS, United Kingdom.

^9^ Department of Biology, University of York, York, United Kingdom.

† These authors contributed jointly to this work

Supplementary Methods

**Cell Lines:** From ~100 screened colorectal carcinoma lines we selected preliminary lines SW620, COLO320HSR and COLO741 due to low endogenous EGFR expression from microarray data(23) (fig. S1) before focusing on SW620 (COLO741 was found to be a melanoma and COLO320HSR exhibited transfection instability). Lysate protein levels were estimated by radio immunoprecipitation and BCA (Thermo Scientific™ Pierce™). Western blotting was performed using anti-EGFR mouse monoclonal (1:1000, clone 1F4, Cell Signaling Technology®) and anti-β-tubulin mouse monoclonal (1:1000, Sigma-Aldrich®) antibodies prepared in TBS-T, 5% milk, incubated overnight at 4ºC, then incubated with secondary antibody of polyclonal rabbit anti-mouse conjugated to horseradish peroxidase (Dako) at 1:10,000 and 1:100,000 for EGFR and β-tubulin, respectively, prior to chemiluminescence exposure (Amersham Biosciences). Cell lines with intermediate EGFR expression were used as positive controls.

Plasmid perbB1-EGFP-N1 (donated, Philippe Bastiaens) was used for transformations, comprising human *EGFR* insertion into enhanced GFP backbone pEGFPN1 plus kanamycin resistance. 1 day pre-transfection, 200,000 SW620 cells in 1ml growth media were seeded into a 12-well plate, adding 2μg of pEGFR-EGFP (Invitrogen), 200μl Invitrogen Opti-MEM® I Reduced Serum Medium, 1μl of Plus^TM^ Reagent and 6μl of Lipofectamine® LTX to each. DNA-lipid complexes were added dropwise to cells, incubated 5h (5% CO_2_, 37 ºC), then media exchanged to normal the following day. Cells were reseeded onto 15cm plates in Gibco® Dulbecco’s modified eagle media (DMEM) supplemented with 4.5g/l glucose, pyruvate, L-glutamine and phenol red plus 2μg/ml Gibco™ Geneticin® (G418 sulfate). Colonies were isolated using a silicon cloning cylinder (Corning®), harvested by trypsinization and transferred in a 12-well plate. Flow cytometry based fluorescence assisted cell sorting (FACS) was used to sort cells into three fractions on the basis of fluorescence intensity using the lowest intensity fraction (i.e. lowest EGFR-GFP expression) for subsequent experiments to minimize effects of expression variability across the cell population. Transgene expression was confirmed by imaging live and immunofluorescently stained fixed cells with confocal, and western blotting. For stimulation with EGF we used an EGF concentration (either unlabeled or as EGF-RMR in dual-color TIRF microscopy) equivalent to 100ng/ml. This level resulted in clear phosphorylation activity on western blots. It is consistent with high physiological levels found in prostate and breast tissue – in particular 50-500ng/mL found in high EGF bodily fluids including prostate fluid (45) or 30-300ng/mL found in breast cysts (46).

For CHO-K1, cDNA encoding EGFR tagged with EGFR-Halo was generated by replacing cDNA encoding YFP protein in human EGFR-YFP plasmid (donated, Ivan Nabi), with that of Halo 7-tag protein (Promega), with insertion of a 45-base linker (15 aa, sequence 3SGGG) between EGFR and Halo. cDNA encoding human Erb2 (National Institute of Technology and Evaluation Biological Resource Centre 2-49-10,Nishihara,Shibuya-ku,Tokyo 151-0066 Japan) was fused at its C-terminus with mGFP (EGFP with A206K mutation), placed in the pOSTet153T vector, inserting an 18-base linker (6 aa, with sequence of three SG repeats) between Erb2 and mGFP. All constructs were confirmed by DNA sequencing.

**Total Copy Number:** The number of EGFR-GFP for SW620:EGFR-GFP on the cell surface was estimated by integration(25) of pixel intensities of the cell area in TIRF corrected for autofluorescence using parental SW629 strain. SW620:EGFR-GFP mean pixel fluorescence was calculated for every cell from a region segmented from brightfield using Sobel edge detection, morphologically dilating by a 7 pixel radius disk to minimize cell edge effects. Copy number was estimated by multiplying this value over the area of the cell, approximated as a 14µm diameter sphere.

For tracked foci in TIRF we see a mean stoichiometry of 12.8 EGFR-GFP with 850 molecules tracked in foci per cell. The total number of tracks is at least 3x this as we detect only the basal membrane which we estimate to be 2,520 EGFR molecules associated into foci per cell in total, indicating ≤197,480 molecules per cell not directly detected in TIRF. If these ‘pool’ foci have a maximum theoretical stoichiometry of S(max), copy number C=197,480 molecules per cell, mean area A=615μm^2^ then to fail to detect separate foci as distinct on the basis of optical resolution w=0.23μm the separation of nearest neighbor EGFR foci ≤w. At a separation of w the equivalent average area occupied by each focus on the membrane is π(w/2)^2^. Therefore A=Cπ(w/2)^2^/S, which suggests S(max) is approximately 40 molecules.

**Mobility.** 2D mean square displacements (MSDs) of tracks was calculated in MATLAB from the centroid at time *t*, (*x*(*t*),y(*t*)), assuming *N* consecutive image frames, and time interval τ=*n*Δ*t*, where *n* is a positive integer with Δ*t* frame integration time(47):

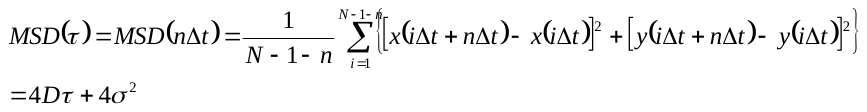

The localization precision is *σ*=40*±*20nm. The apparent diffusion coefficient *D* was estimated from linear fits to the first three points in MSD *vs.* τ, constrained to pass through 4*σ*^2^ at τ = 0, allowing *σ* to vary 20‑60nm.

**Colocalization.** For SW620:EGFR-GFP, colocalization between EGFR-GFP and EGF-TMR was calculated as the overlap integral between green/red foci in MATLAB, whose centroids were within 5 pixels(27). Assuming two normalized, 2D Gaussian intensity distributions $g_{1}(x,y)$ and $g_{2}(x,y)$, centered around (*x*_1_, *y*_1_) with width *σ*_1_, and centered around (*x*_2_, *y*_2_) with width *σ*_2_ for green/red foci respectively, the overlap integral *ν* is(27):

$$v=exp(-\frac{\Delta r^{2}}{2\left( \sigma_{1}^{2}+\sigma_{2}^{2} \right)})$$

where:

$\Delta r^{2}= \left( x_{1}-x_{2} \right)^{2}+{(y_{1}-y_{2})}^{2}$.

Our simulations indicated green/red foci pairs with identical centroids have an overlap of ~0.75, therefore we used a 0.75 threshold for colocalization detection.

For dwell time analysis in CHO-K1 we used a similar method(48, 49), detecting colocalized green/red foci pairs then measuring the duration over which this separation remained ≤180 nm, and generated a histogram distribution of these that could be fitted by the sum of two exponential decay functions with time constants t_1_=53±2ms and t_2_=335±100ms. To determine the time constant associated with random colocalization we rotated the red images by 180° prior to performing the same analysis, indicating a dwell time distribution fitted with a single exponential function t_rot_=52±2 ms, identical within error to t_1_, suggesting t_2_  is associated with non-random colocalization.

**Modelling foci and overlap probability.** To test that foci remained detectable in the higher noise *in vivo* environment, we simulated foci over a 10 molecule range of stoichiometry using the same noise conditions as measured from real cell images. Foci were simulated as 2D Gaussians with the same characteristic intensity as *in vitro* and *in vivo.* Ten images of ten foci were simulated for each stoichiometry and the number of false postives and negatives plotted (fig.5A). The probability that ≥2 fluorescent foci are separated by less than the optical resolution was determined in MATLAB using a previous model(27) at experimental foci surface densities. The apparent stoichiometry distribution from overlapping foci was modelled by convolving a Poisson distribution generated from the probability of overlap with the expected intensity distribution of an isolated multimer. The latter was obtained by scaling the width of the single fluorophore intensity distribution (fig. S5) by *S*^1/2^ where *S* is the model stoichiometry. A random Monte Carlo stoichiometry was generated from a population distribution of oligomeric EGFR with stoichiometry sampled from a Poisson distribution with mean equal to the most likely observed experimental stoichiometry (6 molecules) resulting in a goodness-of-fit *R*^2^=0.4923.

**Statistics.** Two-tailed Student’s *t-*tests were performed for comparisons between pairs of datasets to test null hypothesis that data in each was sampled from the same statistical distribution assuming (n_1_+n_2_-2) degrees of freedom where n_1_ and n_2_ are the number of data points in each distribution and by convention that *t* statistic values which have a probability of confidence P>0.05 are statistically not significant. For TIRF each cell was defined as a biological replicate sampled from the cell population with sample sizes of 10-117 cells per condition. Technical replicates are not possible with irreversible photobleaching, nevertheless. Differences between colocalization dwell times were assessed using the Brunner-Munzel rank order test.

**Kinetic modelling of EGFR ligand binding, dimerization, endocytosis and recycling**

The time-dependent kinetic model that we have developed considers: (i) that EGF ligand can bind to both EGFR receptor monomers and dimers, (ii) that ligated EGFR monomers and singly- or doubly-ligated dimers can be internalized via endocytosis, and (iii) that endocytosed receptors can be recycled back to the plasma membrane. EGFR degradation is not included in the model, as the half-life of EGFR degradation after EGF activation is typically of the order of hours(29) and hence has a negligible impact considering the time scale of our measurements (~40min).

In what follows, we use the following notation: R for receptor monomers, RR for receptor dimers, L for EGF ligand, RL for ligated receptor monomers, RRL for singly-ligated receptor dimers, RRL2 for doubly-ligated receptor dimers, RL^inside^ for endocytosed RL, RRL^inside^ for endocytosed RRL and RRL2^inside^ for endocytosed RRL2. Our model considers the following reversible reactions for ligand binding and receptor dimerization:

R + L → RL with on-rate constant k_11,on_, off-rate constant k_11,off_ and equilibrium association constant $K_{11}$ = k_11,on_**/**k_11,off_;

RR + L → RRL with on-rate constant k_21,on_, off-rate constant k_21,off_ and equilibrium association constant $K_{21}$ = k_21,on_**/**k_21,off_;

RRL + L → RRL2 with on-rate constant k_22,on_, off-rate constant k_22,off_ and equilibrium association constant $K_{22}$ = k_22,on_**/**k_22,off_;

R + R → RR with on-rate constant l_20,on_, off-rate constant l_20,off_ and equilibrium dimerization constant $L_{20}$ = l_20,on_**/**l_20,off_;

RL + R → RRL with on-rate constant l_21,on_, off-rate constant l_21,off_ and equilibrium dimerization constant $L_{21}$ = l_21,on_**/**l_21,off_,;

RL + RL → RRL2 with on-rate constant l_22,on_, off-rate constant l_22,off_ and equilibrium dimerization constant $L_{22}$ = l_22,on_**/**l_22,off_.

Additionally, we have the following non-reversible endocytosis and recycling reactions:

RL → RL^inside^ → R

RRL → RRL^inside^ → RR

RRL2 → RRL2^inside^ → RR.

For the last three reactions, the first arrow (endocytosis) is considered to have a rate k_endoc_ and the second arrow (recycling back to the plasma membrane) is considered to have a rate k_recycle_. We consider only endocytosis of ligated receptors (RL, RRL, RRL2) and, in the first instance, we assume that recycling returns unligated receptors to the plasma membrane. We assume the same endocytosis and recycling rates for the three reactions above.

Considering all the above reactions, it is possible to write a system of simultaneous rate equations as follows:

$\frac{d[\mathrm{RR}]}{dt}$= l_20,on_ [R]^2^- l_20,off_ [RR] - k_21,on_[RR][L]+k_21,off_[RRL] + k_recycle_[RRL^inside^] + k_recycle_[RRL2^inside^],

$\frac{d[\mathrm{RL}]}{dt}$= k_11,on_[R][L] - k_11,off_[RL] - k_endoc_[RL] - l_21,on_[RL][R] + l_21,off_[RRL] - l_22,on_ [RL]^2^ + l_22,off_[RRL2],

$\frac{d[\mathrm{RRL}]}{dt}$= k_21,on_[RR][L] - k_21,off_[RRL] - k_endoc_[RRL] + l_21,on_[RL][R] - l_21,off_[RRL] - k_22,on_[RRL][L] + k_22,off_[RRL2],

$\frac{d[RRL2]}{dt}$= k_22,on_[RRL][L] - k_22,off_[RRL2] - k_endoc_[RRL2] + l_22,on_ [RL]^2^ - l_22,off_[RRL2],

$\frac{d[\mathrm{RL}^{\mathrm{inside}}]}{dt}$= k_endoc_[RL] - k_recycle_[RL^inside^],

$\frac{d[\mathrm{RRL}^{\mathrm{inside}}]}{dt}$ = k_endoc_[RRL] - k_recycle_[RRL^inside^],

$\frac{d[{RRL2}^{\mathrm{inside}}]}{dt}$ = k_endoc_[RRL2] - k_recycle_[RRL2^inside^],

[R] = R_total_ – ([RL] + [RR] + [RRL] + [RRL2] + [RL^inside^] + [RRL^inside^] + [RRL2^inside^]).

Here t is time and the square brackets indicate concentration of reactant/product, with all concentrations being functions of time, and all rates defined as above. R_total_ is the total concentration of receptors (including both receptors on the surface and internalised), i.e. the total number of receptor molecules per cell. This value is set to 200,000-400,000 molecules/cell (depending on the model parameters used) in order to match our observations of approximately 200,000 receptors on the cell surface in experiments.

In order to solve the system of equations we consider the following initial conditions: [RL](t=0) = [RRL](t=0) = [RRL2](t=0) = [RL^inside^](t=0) = [RRL^inside^](t=0) = [RRL2^inside^](t=0) = 0, i.e., that all ligated and internalised components are zero at t=0, and we set the initial concentrations of unligated receptor monomers and dimers to certain fractions of R_total_, typically ~5% monomer fraction and ~95% dimer fraction, in agreement with the fact that we observe mostly clusters of receptors and hardly any monomers in our experiments. The ligand concentration, [L], used is that in our experiments: 100 ng/ml ~ 15.6 nM.

The rate constants for monomer ligand binding/unbinding are given values $k_{11,on}={10}^{6} M^{-1}s^{-1}$ and $k_{11,off}={10}^{-3} s^{-1}$, so that the equilibrium association constant is $K_{11}={10}^{9} M^{-1}$. These rate constants are similar to those previously reported for EGFR monomers(50) and ensure that the time scale of equilibration of receptor binding is ~5-10 min, in agreement with measurements reported in live cells at 37°C (50, 51) . We assume that the ligand binding/unbinding rate constants for unligated receptor dimers are the same as for unligated monomers, i.e., $k_{21,on}=k_{11,on}$ and $k_{21,off}=k_{11,off}$ (hence $K_{21}=K_{11}$). This assumption is motivated by the fact that the equilibrium association constants for R and RR binding to ligand for EGFR have been reported to be approximately the same, i.e., $K_{21}\approx K_{11}$, as obtained from measurements in cells at 4°C(22). The same study reported that the equilibrium association constant for ligand binding to singly-ligated receptor dimers is ~14 times lower than that for unligated monomers/dimers, i.e., $K_{22}\approx K_{11}/14$. To include this reported lower affinity of ligand for singly-ligated receptors, we choose on/off-rate constants for ligand binding to RRL of $k_{22,on}=k_{11,on}/7$ and $k_{22,off}={2\times k}_{11,off}$. Note that these conditions imply that there is negative cooperativity of EGFRs in ligand binding, as $K_{22}<K_{11}/4$.

As for receptor dimerization, equilibration is achieved in shorter times, of the order of ~0.1-1s, as measured for G-protein-coupled receptors (GPCRs) in living cells at 37°C (52). For our model, we choose an equilibrium dimerization constant $L_{20}={10}^{-3}{(molecules/cell)}^{-1}$ in units of 2D concentration (with 1 receptor molecule per ${\mu m}^{2}$ corresponding to approximately 3,000 receptors/cell), 10 times higher than the value measured for GPCRs $({10}^{-4}{(molecules/\mathrm{cell})}^{-1}$). We use $l_{20,on}={10}^{-4} {(molecules/cell)}^{-1}s^{-1}$ and $l_{20,off}={10}^{-1} s^{-1}$ to guarantee that the time scale of dimer formation equilibration is correct and to ensure that there are very few monomers on the cell surface at equilibrium (~5% monomers, ~95% dimers), in agreement with our own experimental observations. For the dimerization rate of RL and R, we use $l_{21,on}=l_{20,on}$ and $l_{21,off}=l_{20,off}$ as the respective equilibrium dimerization constants reported from measurements in cells at 4°C are very similar ($L_{21}\approx L_{20}$) (22). As the dimerization rate $L_{22}$ for RL+RL was reported to be ~10 times lower than $L_{20}$ in the same study, we choose as model parameters $l_{22,on}=l_{20,on}/5$ and $l_{22,off}={2\times l}_{20,off}.$

We set the rate of endocytosis of ligated receptors in our model to $k_{\mathrm{endoc}}=6\%/\min=0.06/60s^{-1}$, in agreement with previous reports of EGFR endocytosis rates in living cells at 37°C in the presence of high EGF concentrations similar to that in our experiments, which are in the range $3-10\%/\min$ (22) (50). EGFR endocytosis rates at low ligand concentrations (near physiological levels [EGF]~1ng/ml) are typically in the range $15-30\%/\min$. At higher EGF concentrations (~100 ng/ml) and high receptor numbers (as in our experiments), the clathrin-endocytosis pathway saturates, and the endocytosis rate is lower ($3-10\%/\min$). Endocytosis rates in the absence of EGF ligand are greatly decelerated with respect to the above rates(22). The recycling rate is set to $k_{\mathrm{recycle}}=10\%/\min=0.1/60s^{-1}$ (22).

Using all the above parameter values, we solve the above system of differential equations in order to find [R], [RL], [RR], [RRL], [RRL2], [RL^inside^], [RRL^inside^] and [RRL2^inside^] as a function of time. Additionally, we calculate the fractional saturation on the cell surface, $Y^{\mathrm{surface}}$, equal to the ratio of EGF to EGFR molecules (EGF:EGFR ratio) on the plasma membrane. This is given by:

$Y^{\mathrm{surface}}= \frac{{[R_{\mathrm{bound}}]}^{\mathrm{surface}}}{{[R_{\mathrm{bound}}]}^{\mathrm{surface}}+{[R_{\mathrm{unbound}}]}^{\mathrm{surface}}}=\frac{[\mathrm{RL}]+{[\mathrm{RRL}]}/2+[RRL2]}{[R]+[\mathrm{RL}]+[\mathrm{RR}]+[\mathrm{RRL}]+[RRL2]}$ .

Here, we account for the fact that all concentrations involving receptors in the model are obtained as numbers of receptor molecules per cell. The value of the EGFR:EGF ratio obtained from our experimental measurements is ~4 on average (and ~2 as a modal peak value), which would correspond to a $Y^{\mathrm{surface}}$ value in our model of ~0.25 (and peak value ~0.5).

The calculated variables as a function of time for the above-mentioned parameters are shown in fig. 4A in the main text. We use R_total_ = 310,000 molecules/cell, which is the value that results in a total number of receptor molecules on the cell surface (excluding internalised receptors) at equilibrium of 200,000, in agreement with our observations. We find that the fractional saturation at the cell surface is $Y^{\mathrm{surface}}\sim0.66$, i.e. the EGFR:EGF ratio predicted by the model is ~1.5.

The results for $Y^{\mathrm{surface}}$ (at equilibrium) do not change significantly upon various changes of the model parameters. Changing the value of R_total_ by ±100,000 receptors/cell only alters the result by <2%. Increasing the endocytosis rate to 10%/min yields $Y^{\mathrm{surface}}\sim0.62$ and decreasing it to 3%/min yields $Y^{\mathrm{surface}}\sim0.72$. Increasing the recycling rate up to 40%/min does not change the value of $Y^{\mathrm{surface}}$ at equilibrium; decreasing it to 0 results in $Y^{\mathrm{surface}}\sim0.83$ and a largely reduced number of receptors at the cell surface (~30,000) after 40 minutes. Increasing or decreasing all ligand binding on-rate constants by a factor of 2 results in moderately increased or decreased values at equilibrium, $Y^{\mathrm{surface}}\sim0.77$ or $Y^{\mathrm{surface}}\sim0.55$, respectively. Increasing or decreasing all ligand binding off-rate constants by a factor of 2 results in moderately decreased or increased values, $Y^{\mathrm{surface}}\sim0.61$ or $Y^{\mathrm{surface}}\sim0.70$, respectively. Increasing or decreasing all receptor dimerization on-rate constants by a factor of 2 changes the result only by <3%. Changing the factors in $k_{22,on}$and $k_{22,off}$ that guarantee that $K_{22}=K_{11}/14$, to go from $k_{22,on}=k_{11,on}/7$ and $k_{22,off}={2\times k}_{11,off}$ to the extremes $k_{22,on}=k_{11,on}/14$, $k_{22,off}=k_{11,off}$ or $k_{22,on}=k_{11,on}$, $k_{22,off}={14\times k}_{11,off}$ changes the result by at most 3%. Changing the factors that guarantee that $L_{22}=L_{20}/10$ to the extremes $l_{22,on}=l_{20,on}/10$, $l_{22,off}=l_{20,off}$ and $l_{22,on}=l_{20,on}$, $l_{22,off}={10\times l}_{20,off}$, does not change the result at all. Reducing all dimerization on/off-rate constants by a factor of 100 (which increases the dimerization equilibration time from ~1s to ~1min) only increases $Y^{\mathrm{surface}}$ to 0.68 (by 3%). Changing the initial fractions of receptor monomers and dimers at t=0 does not change the results as the equilibration time of dimerization is short (~1s).

Setting the binding on-rate constant for singly-ligated dimers to zero ($k_{22,on}=0$), i.e. assuming extreme negative cooperativity, the result remains the same ($Y^{\mathrm{surface}}\sim0.66$). Setting the binding on-rate constant for unligated dimers to zero ($k_{21,on}=0$) makes a significant difference and results in $Y^{\mathrm{surface}}\sim0.24$, i.e., in an EGFR:EGF ratio at equilibrium on the cell surface of ~4 (see fig. 4B in the main text). This is the case regardless of whether $k_{22,on}$is also set to zero or not, i.e., this result is consistent also with extreme preferential ligand binding to monomers (ligand can bind to receptor monomers but not to dimers). Setting the binding on-rate constant for receptor monomers to zero ($k_{11,on}=0$), i.e. assuming extreme preferential ligand binding to dimers, yields $Y^{\mathrm{surface}}\sim0.63$. In the case of positive cooperativity, setting $k_{22,on}=k_{11,on}$ and $k_{22,off}=k_{11,off}$, the result increases to $Y^{\mathrm{surface}}\sim0.69$. Considering more significant positive cooperativity with $k_{22,on}=10 k_{11,on}$ and $k_{22,off}=k_{11,off}$, we obtain an even higher value, $Y^{\mathrm{surface}}\sim0.82$. Hence, positive cooperativity leads to higher values of $Y^{\mathrm{surface}}$, while negative cooperativity leads to lower values. Hence, the latter agrees better with our experimental results that correspond to an EGFR:EGF ratio ~4 on average (i.e., to $Y^{\mathrm{surface}}\sim1/4$). Model predictions in the case of preferential ligand binding to monomers would agree with experimental observations.

We also tried setting some of the dimerization rate constants to zero as follows. Assuming that R monomers do not dimerize to form RR dimers ($l_{20,on}=0$), we obtain $Y^{\mathrm{surface}}\sim0.49$ and $Y^{\mathrm{surface}}\sim0.69$ if we also set the off-rate constant to zero ($l_{20,off}=0$). Assuming that RL and R do not dimerize to form RRL ($l_{21,on}=0$), we obtain $Y^{\mathrm{surface}}\sim0.92$ and $Y^{\mathrm{surface}}\sim0.69$ if we also set the off-rate constant to zero ($l_{21,off}=0$). Hence, a decreased hetero-dimerization rate constant (ligated-unligated) leads to model predictions that are further from our experimental observations. Assuming that RL complexes do not dimerize to form RRL2 ($l_{22,on}=0$), we obtain $Y^{\mathrm{surface}}\sim0.53$ and $Y^{\mathrm{surface}}\sim0.68$ if we also set the off-rate to zero ($l_{22,off}=0$). Setting both R and RL homo-dimerization rate constants (unligated-unligated and ligated-ligated) to zero ($l_{20,on}=0$ and $l_{22,on}=0$), the model yields $Y^{\mathrm{surface}}\sim0.47$. Hence, a reduced homo-dimerization on-rate constant for unligated and/or ligated monomers (i.e. a reduced $l_{20,on}$ and/or a reduced $l_{22,on}$) yields model results that are closer to our experimental observations (a lower $Y^{\mathrm{surface}}$ value).

We also tested in our model ligand binding and dimerization rate constants that result in the same equilibrium association constants as those reported from measurements in cells at 4°C, i.e., $K_{11}\approx5\times{10}^{9}M^{-1}$, $K_{21}\approx K_{11}$, $K_{22}\approx K_{11}/14$, $L_{20}\approx2\times{10}^{-5}{(molecs/cell)}^{-1}$, $L_{21}\approx L_{20}$ and $L_{22}\approx L_{20}/10$ (22). In this case, we set the model parameters to R_total_ = 310000 molecules/cell, 5% and 95% initial fractions of monomers and dimers respectively, $\left[ L \right]=15.6\times{10}^{-9}M$, $k_{\mathrm{endoc}}=0.06/60s^{-1}$, $k_{\mathrm{recycle}}=0.1/60s^{-1}$ (up to here all parameters are the same as previously), and $k_{11,on}={10}^{7} M^{-1}s^{-1}$, $k_{11,off}={2\times10}^{-3} s^{-1}$, $k_{21,on}=k_{11,on}$, $k_{21,off}=k_{11,off}$, $k_{22,on}=k_{11,on}/7$ and $k_{22,off}={2\times k}_{11,off}$, $l_{20,on}\approx2\times{10}^{-5}{(molecs/cell)}^{-1}s^{-1}$ and $l_{20,off}=1 s^{-1}$, $l_{21,on}=l_{20,on}$, $l_{21,off}=l_{20,off}$, $l_{22,on}=l_{20,on}/5$ and $l_{22,off}={2\times l}_{20,off}.$ Using these parameters, the value of the surface fractional saturation (EGF:EGFR ratio) is $Y^{\mathrm{surface}}\sim0.96$ (see Figure 4C in the main text). The result remains the same if all binding on/off-rate constants are multiplied or divided by 10, or if all dimerization on/off-rate constants are multiplied or divided by 10, keeping the equilibrium association constants the same. This value of $Y^{\mathrm{surface}}\sim0.96$ close to 1 is consistent with the results for our ligand concentration from alternative models based on equilibrium equations (that do not include time dependency)(22). It is important to note that the on/off-rate constants can have a strong temperature dependence and that results measured at 4°C can differ significantly (up to a factor of 10-100) from measurements at 37°C (53).

In conclusion, the value we obtain experimentally for the EGFR:EGF ratio in living cells at 37°C agrees better with model predictions in the cases in which there is negative cooperativity for ligand binding and preferential ligand binding to monomers or reduced homo-dimerization on-rate constants (for unligated-unligated and ligated-ligated dimerization).

Additionally, we considered a model in which ligated receptors are recycled back to the plasma membrane ligated, as opposed to unligated. In this case, the system of rate equations that we solve is the following:

$\frac{d[\mathrm{RR}]}{dt}$= l_20,on_ [R]^2^- l_20,off_ [RR] - k_21,on_[RR][L]+k_21,off_[RRL],

$\frac{d[\mathrm{RL}]}{dt}$= k_11,on_[R][L] - k_11,off_[RL] - k_endoc_[RL] - l_21,on_[RL][R] + l_21,off_[RRL] - l_22,on_ [RL]^2^ + l_22,off_[RRL2] + k_recycle_[RL^inside^],

$\frac{d[\mathrm{RRL}]}{dt}$= k_21,on_[RR][L] - k_21,off_[RRL] - k_endoc_[RRL] + l_21,on_[RL][R] - l_21,off_[RRL] - k_22,on_[RRL][L] + k_22,off_[RRL2] + k_recycle_[RRL^inside^],

$\frac{d[RRL2]}{dt}$= k_22,on_[RRL][L] - k_22,off_[RRL2] - k_endoc_[RRL2] + l_22,on_ [RL]^2^ - l_22,off_[RRL2] + k_recycle_[RRL2^inside^],

$\frac{d[\mathrm{RL}^{\mathrm{inside}}]}{dt}$= k_endoc_[RL] - k_recycle_[RL^inside^],

$\frac{d[\mathrm{RRL}^{\mathrm{inside}}]}{dt}$ = k_endoc_[RRL] - k_recycle_[RRL^inside^],

$\frac{d[{RRL2}^{\mathrm{inside}}]}{dt}$ = k_endoc_[RRL2] - k_recycle_[RRL2^inside^],

[R] = R_total_ – ([RL] + [RR] + [RRL] + [RRL2] + [RL^inside^] + [RRL^inside^] + [RRL2^inside^]).

The solution we obtain with the original set of parameters (R_total_ = 310000 molecs/cell, 5% and 95% initial fractions of monomers and dimers respectively, $\left[ L \right]=15.6\times{10}^{-9}M$, $k_{\mathrm{endoc}}=0.06/60s^{-1}$, $k_{\mathrm{recycle}}=0.1/60s^{-1}$, $k_{11,on}={10}^{6} M^{-1}s^{-1}$, $k_{11,off}={10}^{-3} s^{-1}$, $k_{21,on}=k_{11,on}$, $k_{21,off}=k_{11,off}$, $k_{22,on}=k_{11,on}/7$ and $k_{22,off}={2\times k}_{11,off}$, $l_{20,on}\approx{10}^{-4}{(molecs/cell)}^{-1}s^{-1}$, $l_{20,off}=0.1 s^{-1}$, $l_{21,on}=l_{20,on}$, $l_{21,off}=l_{20,off}$, $l_{22,on}=l_{20,on}/5$ and $l_{22,off}={2\times l}_{20,off}$) is $Y^{\mathrm{surface}}\sim0.81$. Similarly to what occurred for the previously mentioned model predictions for experiments at 37°C, the value of the surface fractional saturation decreases significantly when setting $k_{21,on}$ to zero, to $Y^{\mathrm{surface}}\sim0.36$. Variations upon changes of parameters are very similar to those for the previous model at 37°C. Results from this second model are somewhat further from our experimental observations but the conclusions from above remain valid for both types of recycling processes considered (ligated receptors being endocytosed and recycled back to the plasma membrane unligated or ligated).

**EGFR ligand expression data**

Microarray expression data were generated by a service provided by the Patterson Laboratory in Manchester, UK. . The gene expression data for 78 unique, non-duplicate (not sourced from same patients) colorectal cancer cell lines were obtained by performing microarray using the Affymetrix GeneChip HG-U133 Plus 2.0 microarray. Data is normalized using RMA and batch-removed using Partek Genomics Suite software. These were compared against publicly available RNA-Seq data(54). For microarray, we developed an algorithmic pipeline with the goal of identifying subpopulations within a given data through a statistical machine-learning method. With this pipeline we statistically determined the expressing and non-expressing subpopulations in our 78 CRC microarray cell line data. The statistically determined background threshold closely matches with the background expression range of 100-150 microarray counts shown in previous work done by our lab; this therefore defines our background threshold. Expression level below the background threshold is considered as non-expressing.

The results obtained from different probesets for the microarray where available, and for RNAseq, are given below for both the SW620 cell line. In the case of the TGFA ligand we also compared the SW620 output with that of SW480 (a duplicate cell line of SW620 derived from the same patient):

TGFA ligand:

|  | **SW620** | **SW480** | **Category** |
| --- | --- | --- | --- |
| **Micorarray 211258_s_at** | 100 | 81 | Background |
| **Micorarray 205015_s_at** | 218 | 195 | Very Low |
| **Micorarray 205016_at** | 360 | 327 | High |
| **CCLE RNA-Seq (RPKM)** | 6.04 | 11.04 | Very Low |

(**211258_s_at and 205015_s_at)**

Pearson correlation: 0.93

P-value: 4.27e-35

Note: 2 of the 3 probesets of TGFA (211258_s_at and 205015_s_at) correlate strongly with one another with a r-value of 0.93 while either probesets correlate with 205016_at with a r-value of 0.74 and 0.76, respectively. Further exploring SW480, a duplicate cell line of SW620 derived from the same patient, revealed a similar pattern in all three probes. We conclude that 205106_at is an anomaly that does not follow the pattern of the other two probes. Adding onto the expression level observed in CCLE RNA-Seq data of 6.04, we can conclude a consensus very low expression level of TGFA.

ERBB2 ligand:

|  | **SW620** | **Category** |
| --- | --- | --- |
| **Micorarray 216836_s_at** | 200 | Very Low |
| **Micorarray 210930_s_at** | 74 | Background |
| **CCLE RNA-Seq (RPKM)** | 9.07 | Very Low |

(**216836_s_at and 210930_s_at)**

Pearson correlation: 0.50

P-value: 2.53e-06

Note: Both probesets of ERBB2 (216836_s_at and 210930_s_at) do not correlate as well with a r-value of 0.5 thus we score them individually and look at them separately. Adding onto the expression level observed in CCLE RNA-Seq data, we can conclude a consensus low expression level of ERBB2.

HBEGF ligand:

|  | **SW620** | **Category** |
| --- | --- | --- |
| **Micorarray 222076_at** | 41 | Background |
| **Micorarray 244857_at** | 87 | Background |
| **Micorarray 38037_at** | 189 | Very Low |
| **Micorarray 203821_at** | 239 | Very Low |
| **CCLE RNA-Seq (RPKM)** | 1.53 | Very Low |

(**222076_at and 203821_at)**

Pearson correlation: 0.97

P-value: 5.81e-52

Note: 2 of the 4 probesets of HBEGF (222076_at and 203821_at) correlate strongly with an r-value of 0.97 while 38037_at has a poor correlation with either probesets of 0.02 and 0.02, respectively, and 203821_at has a poor correlation with either probesets of 0.11 and 0.13, respectively. Surprisingly, 38037_at and 203821_at also have a poor correlation with an r-value of 0.07, suggesting that the two probesets have unique patterns. We conclude that both 38037_at and 203821_ are anomalies that do not behave in line with the other two probes. Thus, 222076_at and 203821_at is assumed to represent the general expression pattern of HBEGF. Adding onto the expression level observed in CCLE RNA-Seq data, we conclude a consensus low expression level of HBEGF.

AREG ligand:

|  | **SW620** | **Category** |
| --- | --- | --- |
| **Micorarray 205239_at** | 97 | Background |
| **Micorarray 215564_at** | 34 | Background |
| **CCLE RN-Seq (RPKM)** | 0.103 | Background |

(**205239_at and 215564_at)**

Pearson correlation: 0.17

P-value: 0.13

Note: Both probesets of AREG (205239_at and 215564_at) do not correlate well with an r-value of 0.17 thus we score them individually and look at them separately. Adding onto the expression level observed in CCLE RNA-Seq data, we can conclude a consensus background expression level of AREG.

EREG ligand:

|  | **SW620** | **Category** |
| --- | --- | --- |
| **Micorarray 1569583_at** | 59 | Background |
| **Micorarray 205767_at** | 133 | Background |
| **CCLE RNA-Seq (RPKM)** | 0.407 | Background |

(**1569583_at and 205767_at)**

Pearson correlation: 0.81

P-value: 3.77e-19

Note: Both probesets of EREG (1569583_at and 205767_at) correlate strongly with an r-value of 0.81. Adding onto the expression level observed in CCLE RNA-Seq data, we can conclude a consensus background expression level of EREG.

BTC ligand

|  | **SW620** | **Category** |
| --- | --- | --- |
| **Micorarray 207326_at** | 39 | Background |
| **Micorarray 241412_at** | 16 | Background |
| **CCLE RNA-Seq (RPKM)** | 0.127 | Background |

(**207326_at and 241412_at)**

Pearson correlation: 0.87

P-value: 1.50e-25

Note: Both probesets of BTC (207326_at and 241412_at) correlate strongly with an r-value of 0.87. Adding onto the expression level observed in CCLE RNA-Seq data, we conclude a consensus background expression level of BTC.

EGF ligand:

|  | **SW620** | **Category** |
| --- | --- | --- |
| **Micorarray 206254_at** | 26 | Background |
| **CCLE RNA-Seq (RPKM)** | 0.0029 | Background |

Note: EGF has a single probeset with a background expression level. Adding onto the expression level observed in CCLE RNA-Seq data, we can conclude a consensus background expression level of EGF.

EPGN ligand:

|  | **SW620** | **Category** |
| --- | --- | --- |
| **CCLE RNA-Seq (RPKM)** | 0 | Background |

Note: there is no available microarray data for EPGN. The CCLE RNA-Seq data showed a level of 0 thus we can conclude a background expression level of EPGN.

Lastly, a good example of a high expression level in CRC cell lines is the marker CDH1 since it is the signature gene expressed by all epithelial cells. In our microarray data, the average expression level of CDH1 is ~6000 while in the CCLE RNA-Seq data lie around 45-60.

**Supplementary Tables**

| **Biochemical intervention** | | | **EGFR foci stoichiometry,**  **uncolocalized** | | **EGFR foci stoichiometry, colocalized with EGF** | | N cells |
| --- | --- | --- | --- | --- | --- | --- | --- |
| E | C | T | Mean ± s.e.m  (molecules per EGFR focus) | N tracks | Mean ± s.e.m  **(**molecules per EGFR focus) | N tracks |  |
| - | - | - | 12.8±0.4 | 1,250 | X | X | 19 |
| + | - | - | 10.8±0.2 | 4,741 | 31.1±1.1 | 1,969 | 117 |
| - | + | - | 19.9±1.0 | 531 | X | X | 10 |
| - | - | + | 15.3±0.7 | 408 | X | X | 10 |
| + | + | - | 18.8±0.5 | 916 | 51.0±2.1 | 303 | 25 |
| + | - | + | 16.8±0.4 | 1,273 | 44.2±2.4 | 334 | 27 |

**Table S1. Mean EGFR foci stoichiometry.** Number of tracks in total (N tracks) and individual cells (N cells) in datasets indicated. Biochemical interventions for added EGF (E), cetuximab (C), and trastuzumab (T) shown.

**Supplementary Figures**

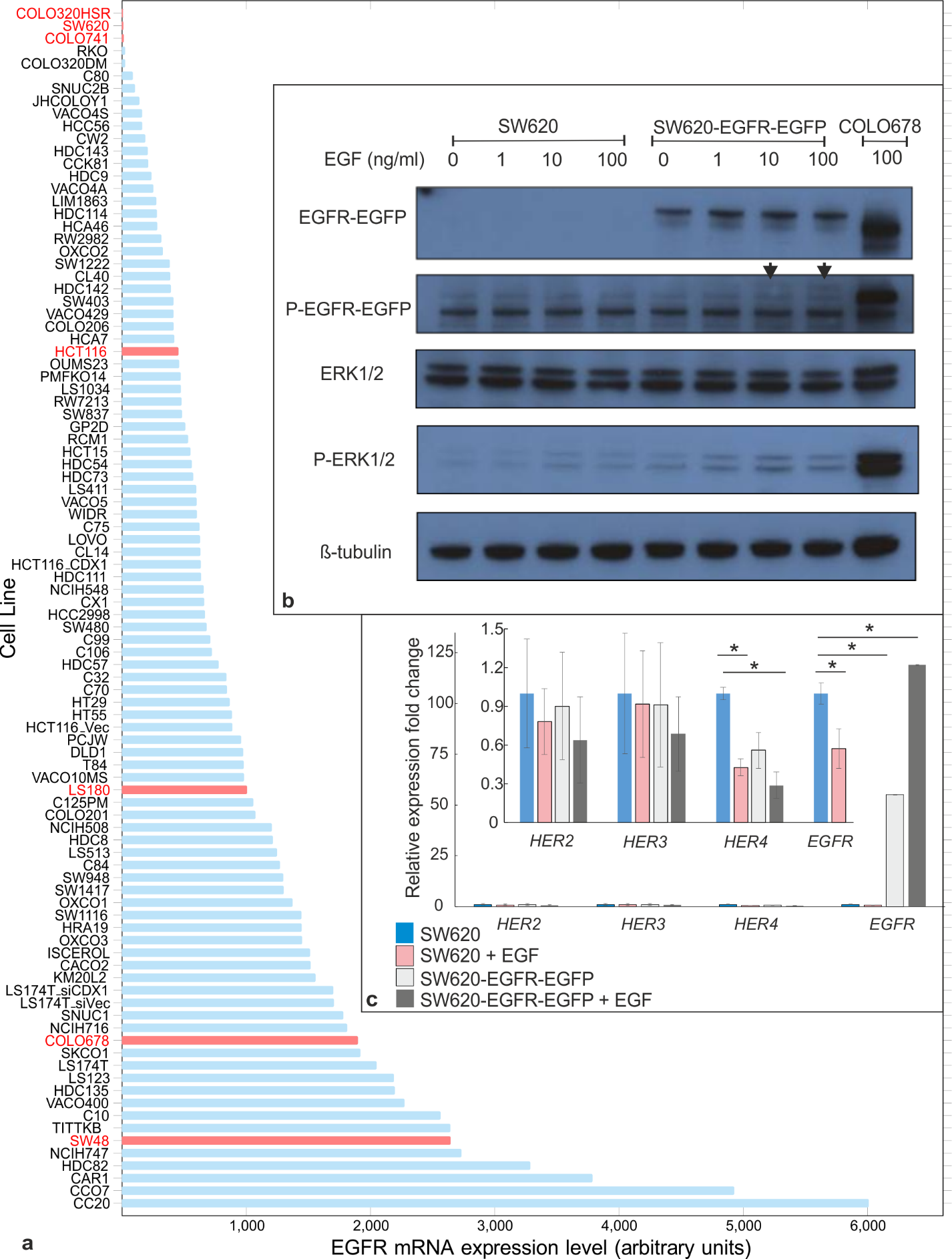

**Fig. S1 EGFR expression levels. A.** The mRNA expression levels were quantified for a colorectal cancer cell line panel using Affymetrix U133+2 mRNA microarray data. Measurements indicated three candidate cell lines, SW620, COLO320HSR and COLO741 (labelled in red, top of panel), as having very low levels of native EGFR expression, as tested in subsequent western blot analysis in comparison to EGFR-expressing cell lines as positive controls (indicated as red columns, middle and bottom of panel). Three candidate cell lines with very low or absent levels of EGFR mRNA (SW620, COLO320HSR, COLO741; Y axis text label in red, top of panel) and a further four positive controls with medium to high levels (HCT116, LS180, COLO678; indicated as red columns, middle and bottom of panel) were selected. Protein levels were confirmed by western blot, as shown for SW620 and COLO678 (fig. 2). **B.** SDS-PAGE performed on protein extracts obtained from SW620 wild type and SW620-EGFR-GFP cells before and after EGF treatment. COLO680 is used as a positive control. SW620 extracts do not have levels of EGFR detectable by SDS-PAGE, unlike SW620-EGFR-GFP, confirming successful transfection and expression of the GFP-tagged EGFR. EGF treatment of the transfected line results at concentrations of 10ng/ml or above results in detectable levels of phosphorylated EGFR (black arrows). EGF treatment of SW620-EGFR-GFP results in increased levels of phosphorylated ERK1/2 kinases, downstream players of the EGFR/ERK pathway, confirming the kinase activity of the receptor. **c.** Fold expression change of *HER2*, *HER3*, *HER4* and *EGFR* relative to *PQLC2* house-keeping gene in SW620 and SW620-EGFR-EGFP cell with or without EGF treatment. Relative fold expression change was calculated based on the ΔΔCt method. The result represents the means and standard errors of four experiments on two biological replicates. *=Student *t*-test P<0.05

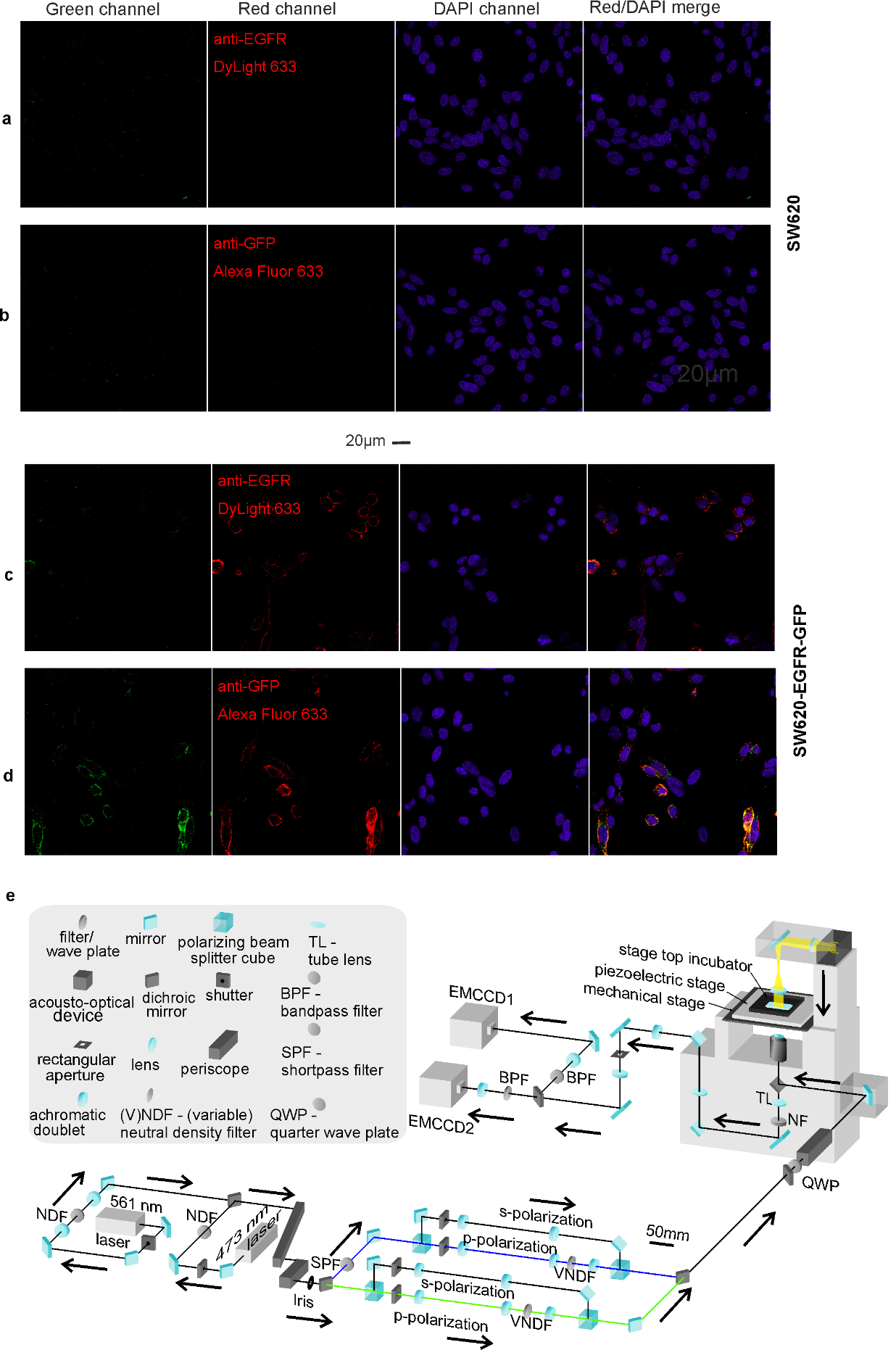

**Fig. S2. Confocal and TIRF characterization.** Confocal microscopy images of fixed cells using GFP, anti-GFP immunofluorescence, and DAPI staining: **(A,B)** non-GFP background cell line SW620; **(C,D)** SW620-EGFR-GFP; **(E)** optical path diagram of bespoke single-molecule TIRF microscope.

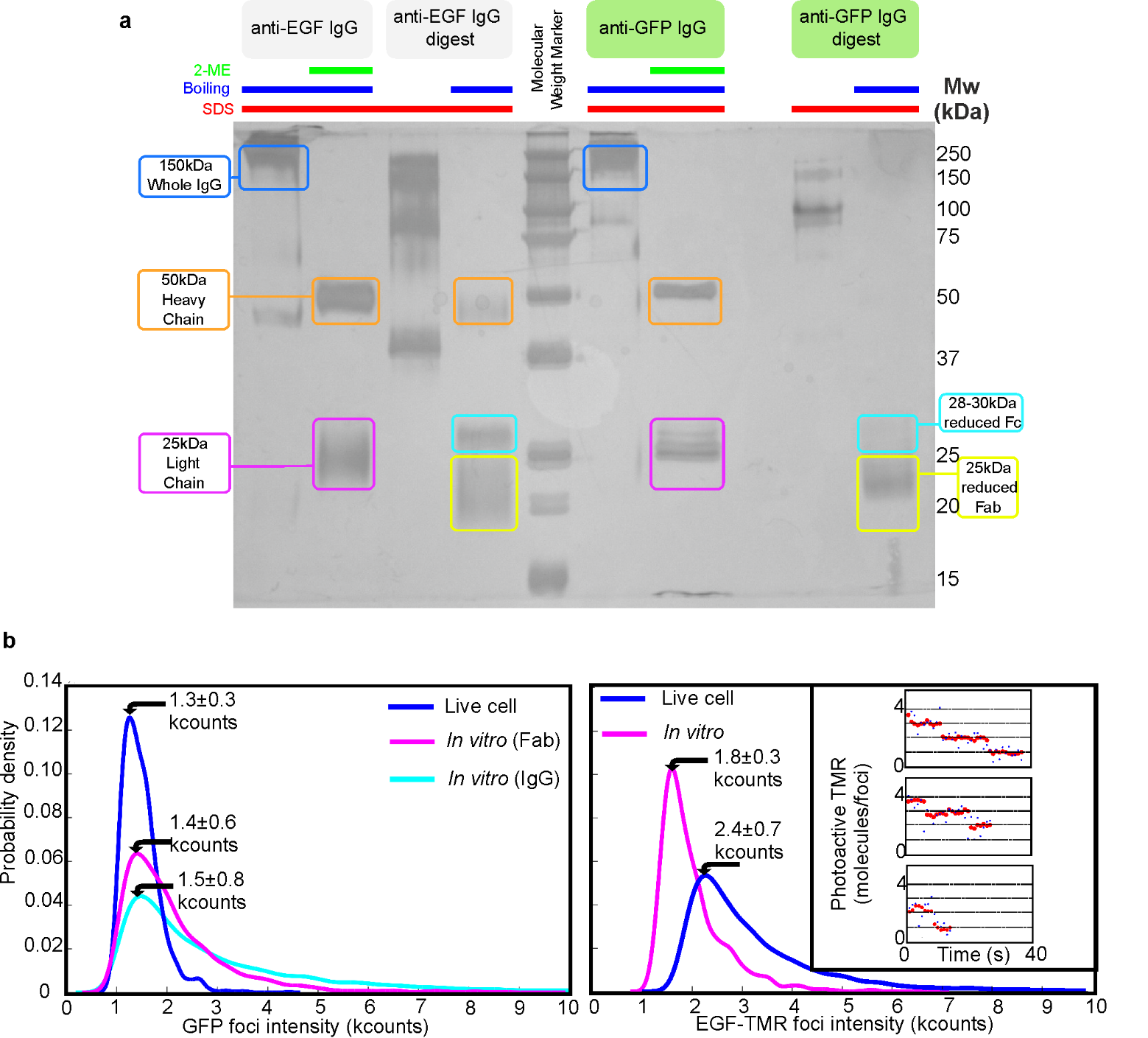

**Fig. S3. Characterization of unitary fluorophore brightness values. (A)** SDS-PAGE gel indicating generation of Fab fragments (yellow) from anti-EGF and anti-GFP IgG antibodies (blue), heavy (orange) and light chains (magenta) indicated with reduced Fc (cyan). **(B)** Kernel density estimation(24) distributions of fluorescent foci intensity values measured in kcounts (i.e. counts x 10^3^) for single GFP (left panel) for live cell, at the end of the photobleach, before EGF is added compared with *in vitro* Fab and whole IgG data. TMR molecule data for *in vitro* EGF-TMR and live cell, at the end of the photobleach, post EGF binding data taken from colocalized EGF-EGFR foci is shown (right panel); inset shows live cell EGF-TMR photobleach steps after EGF has been added, taken from colocalized EGF-EGFR foci, with raw (blue) and Chung-Kennedy filter(55, 56) (red) traces, mean and s.e.m. indicates (arrows).

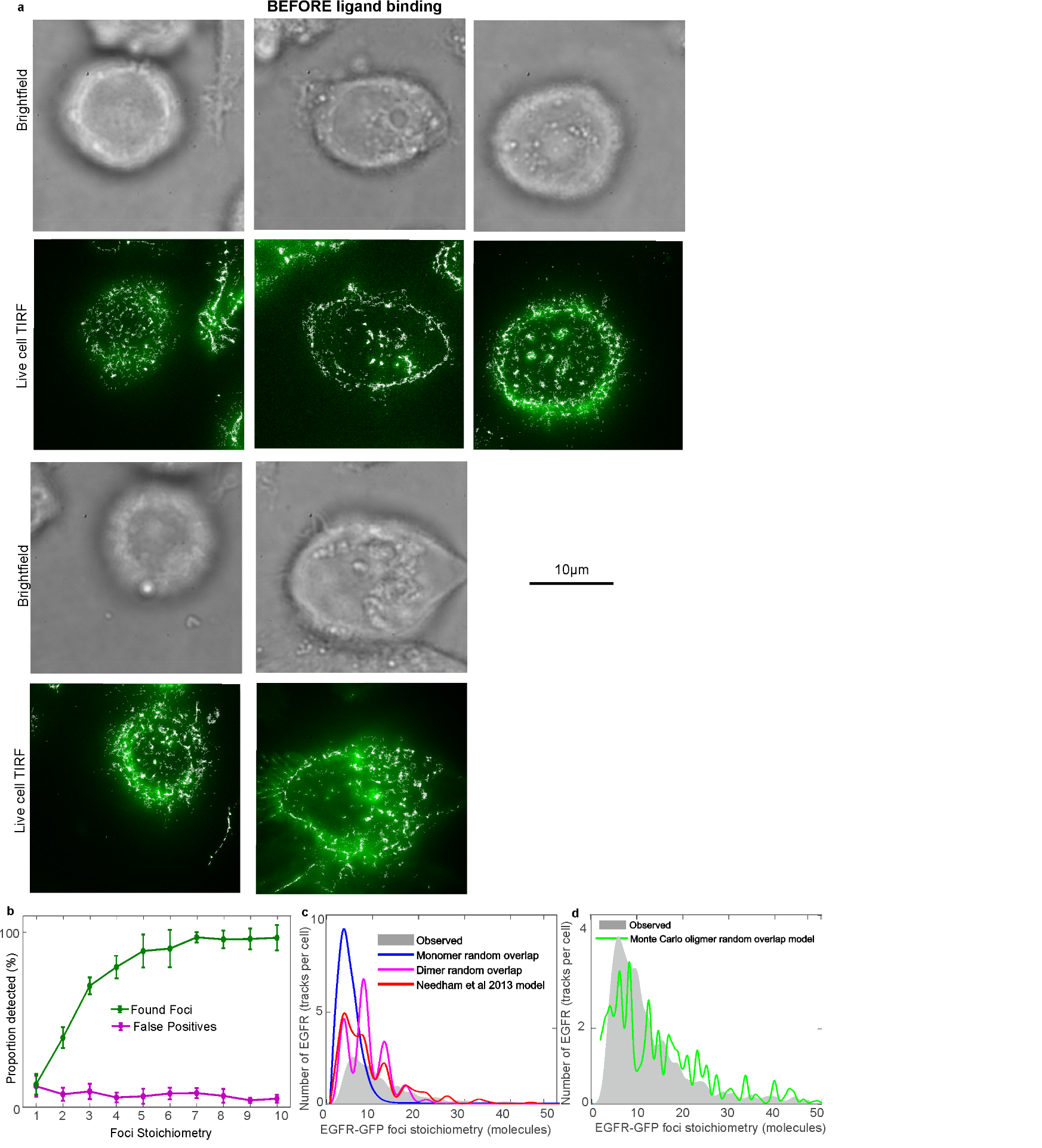

**Fig. S4. More examples of cells before addition of EGF ligand and random foci overlap model. (A)** Brightfield images (grey) and TIRF (green) shown with overlaid foci tracking output (white). We estimate that we detect only *ca.* 13% of available EGFR-GFP in the basal membrane as distinct tracked foci, the remainder contributing to the apparent background resulting in a reduced signal-to-noise ratio than compared to lower expression cell lines. **(B)** Proportion of correctly found and false positive foci in simulations with representative *in vivo* noise. **(C)** Random overlapping foci predictions for **(B)** monomeric (blue) and dimeric EGFR (magenta), and a mixed model oligomer model suggested from a previous single-molecule study (red)(57), all showing poor agreement (*R^2^*<0) to our experimental observations for stoichiometry distribution (grey). **(D)** Monte Carlo Poisson model using an expected average value of 6 molecules for EGFR foci stoichiometry (green) showing a regression fit that can account for *ca.* 50% of the experimental variance (*R^2^*=0.4923) (grey).

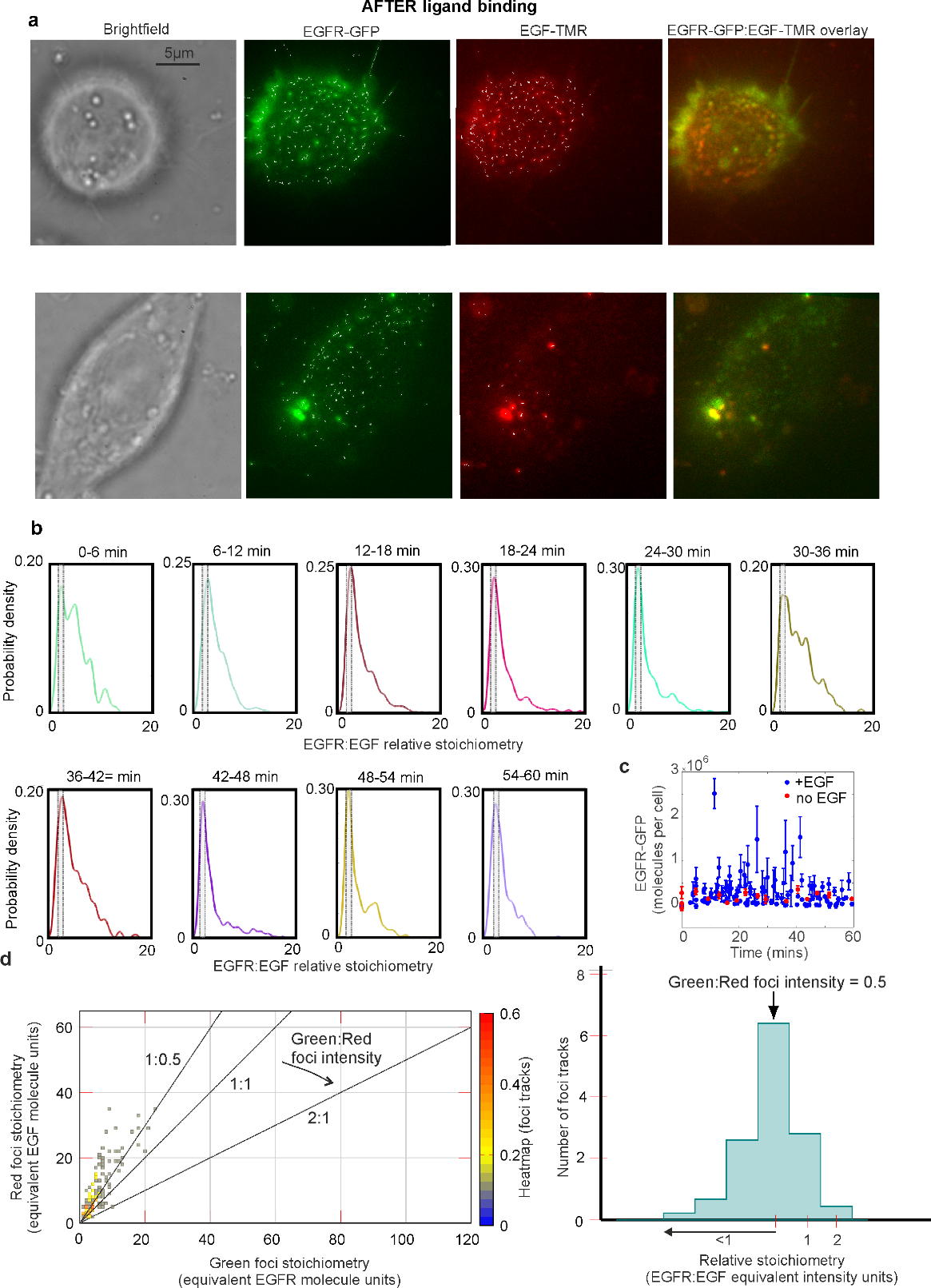

**Fig. S5. Characterizing EGFR and EGF foci stoichiometry after addition of EGF.** **(A)** Two examples of cells taken ~10 min after the addition of EGF: brightfield images (grey), green channel showing EGFR-GFP localization (green), red channel showing EGF-TMR localization (red), and the overlay of green and red channels together (right panels, with yellow indicating regions of high colocalization) are shown here. **(B)** Variation of the EGFR:EGF relative stoichiometry, rendered as kernel density estimations, as a function of incubation time with EGF (shown in 6 min bins). The region corresponding to 2.0 ± 0.5 relative stoichiometry is indicated as a grey rectangle. **(C)** Total cell EGFR-GFP copy number on the plasma membrane cell surface as a function of time, with and without EGF **(D)** Heatmap (left panel) and histogram (right panel) characterizing ‘false’ colocalization due to cellular autofluorescence in green and red channels.

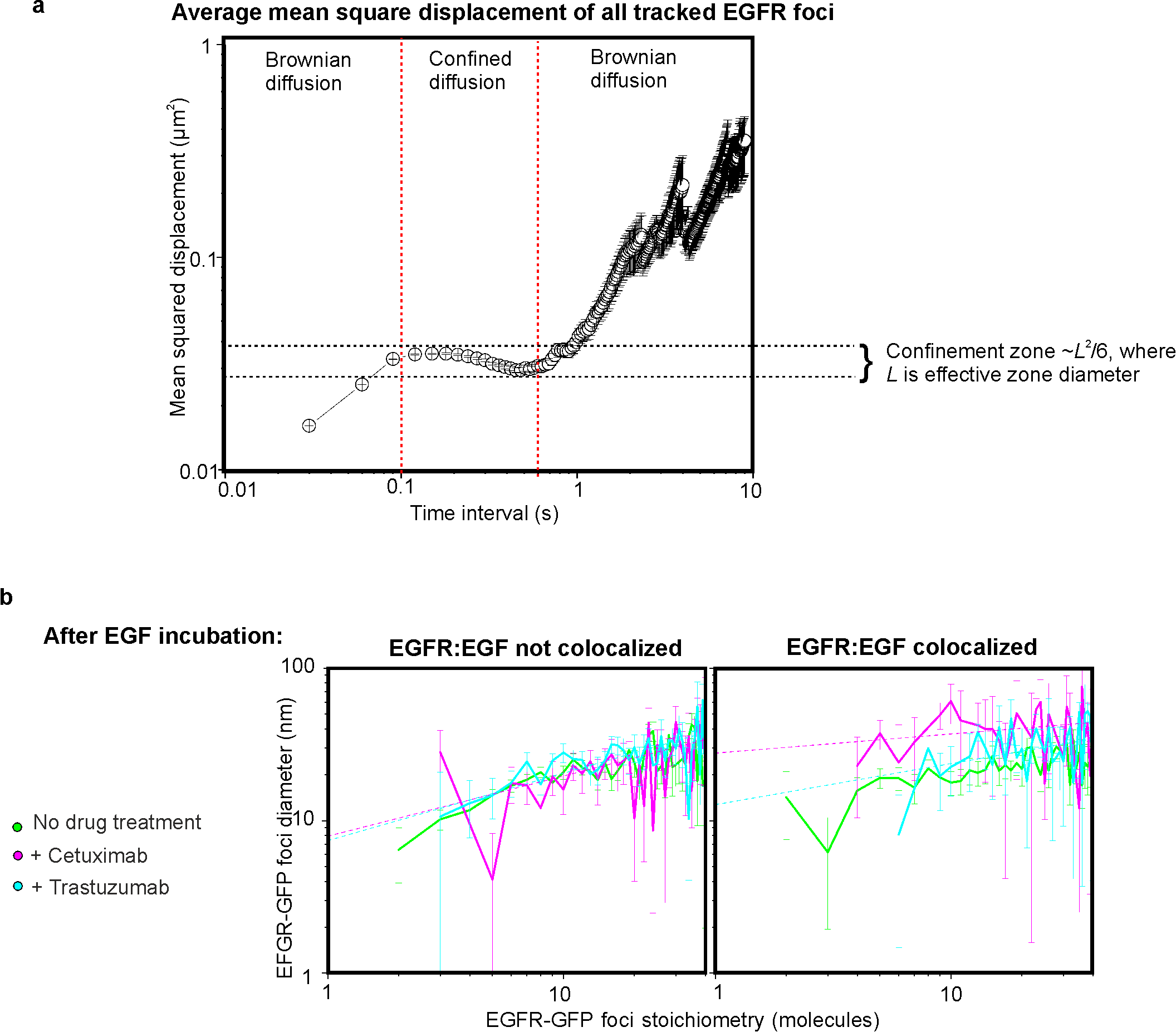

**Fig. S6. EGFR foci diffusion.** (Aa) Log-log plot for average mean square displacement *vs.* time interval for all collated EGFR-GFP foci tracks before addition of EGF, putative confinement zone indicated (dashed lines), from number of foci N=770, acquired from number of cells N=19. (B) Log-log plots of EGFR-GFP foci diameters minus the width for a single GFP molecule *vs.* stoichiometry for not colocalized (left panel) and colocalized foci (right panel), showing cells with no cetuximab or trastuzumab treatment (green, N=6,710 foci, N=117 cells), those treated with cetuximab (magenta, N=1,219 foci, N=25 cells), and those treated with trastuzumab (cyan, N=1,607 foci, N=27 cells), with heuristic power law fit (dash lines), s.e.m. error bars.

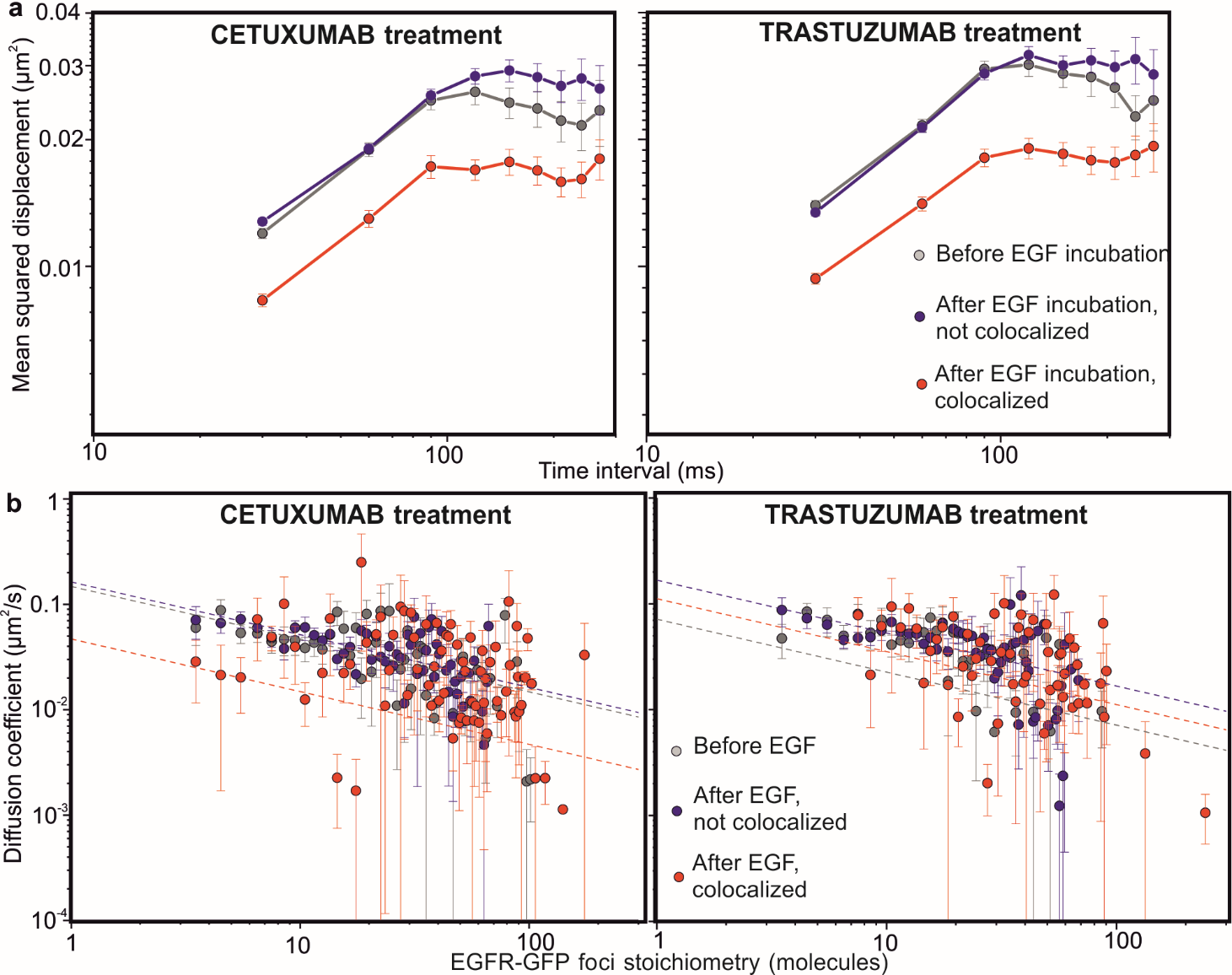

**Fig. S7. Treatment effects on mobility (A)** Log-log plots for average mean squared displacement for time intervals of 300ms or less, and **(B)** log-log plots for variation of apparent microscopic diffusion coefficient *D* with EGFR stoichiometry *S*, fits shown to Stokes-Einstein model assuming *D*~*S*^‑1/2^ (dashed lines) for cetuximab- and trastuzumab-treated cells.

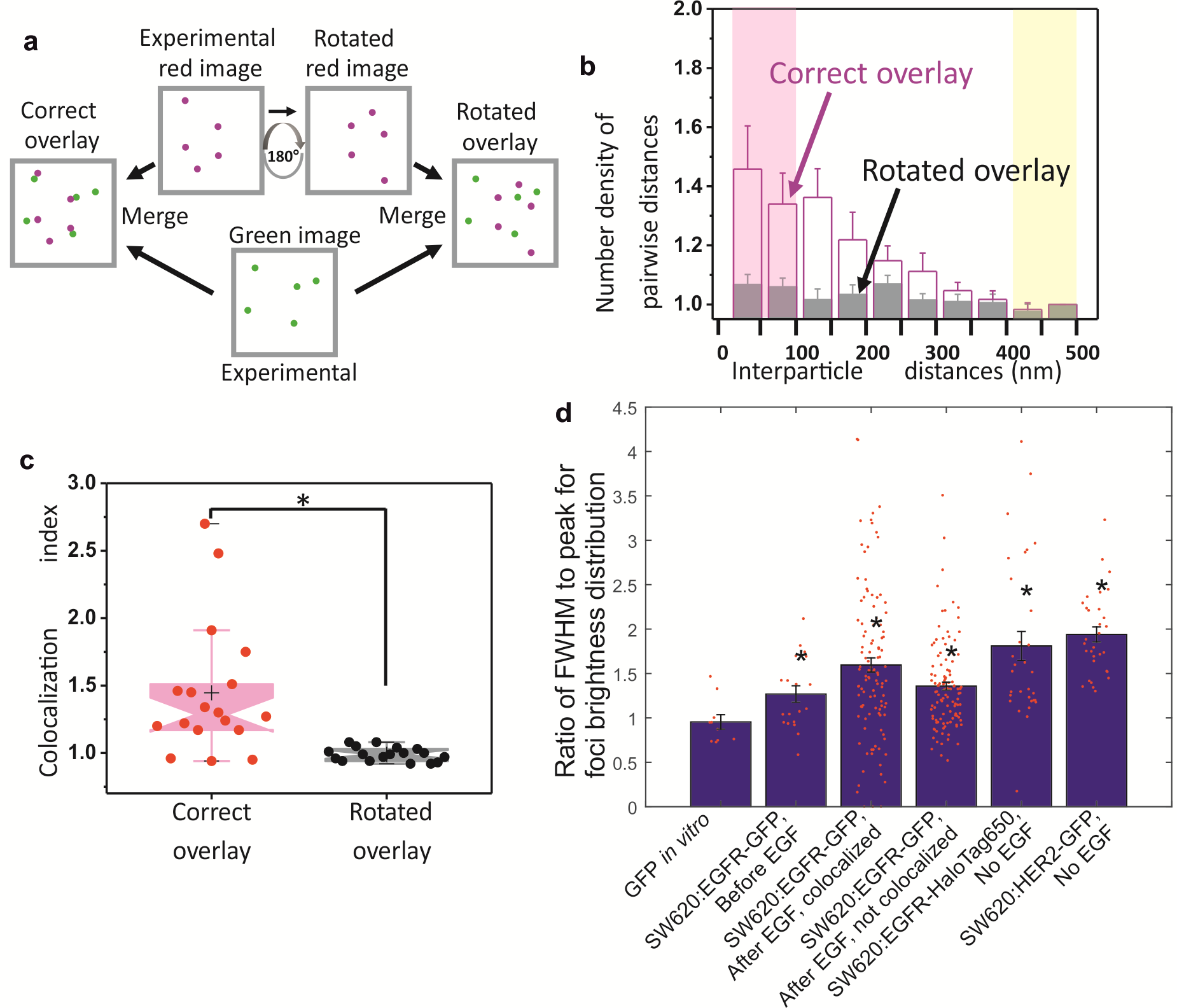

**Fig. S8. EGFR and Her2 colocalisation (A)** In the correct overlay, green and red movies were superimposed. The rotated overlay was used to evaluate random colocalization; the red movie was rotated by 180° and superimposed on the unrotated green movie. **(B)** Distribution of pairwise (interparticle) distances (bin size 50nm) normalized by total number of green-red foci pairs. To parametrize the colocalization level in a simple way we took the ratio of the mean of the number densities of pairwise distances between 0‑100nm (magenta) *vs.* that between 400‑500nm (yellow), termed “Colocalisation index”, s.e.m. error bars, N=20 cells. **(C)** Colocalization indices for “correct” and “rotated” overlays (i.e. with no rotation between green and red channels, and with a 180° rotation for the red channel respectively), showing significant colocalization between EGFR and HER2 (*P* < 0.05, Brunner-Munzel test). Notches, crosses, boxes, and whiskers indicate the median values, mean values, interquartile ranges (25–75%), and the 10–90% ranges, respectively. (D) Comparing clustering between CHO-K1 and SW620 cells. Mean ratios of the full width half maximum (FWHM) divided by peak for the foci intensity distributions of EGFR and HER2 compared with single GFP molecules *in vitro*. Higher ratios than the fluorophore alone are indicative of broader intensity distributions and clustering, s.e.m. error bars over populations of 30-110 cells or 10 fields of view for GFP *in vitro.* Asterix indicates significant differences from GFP *in vitro* below P=0.05 by Wilcoxon-Mann-Whitney test, specific P values =0.020, 0.0039, 0.0030, 0.00040, 7.54E-06 respectively.

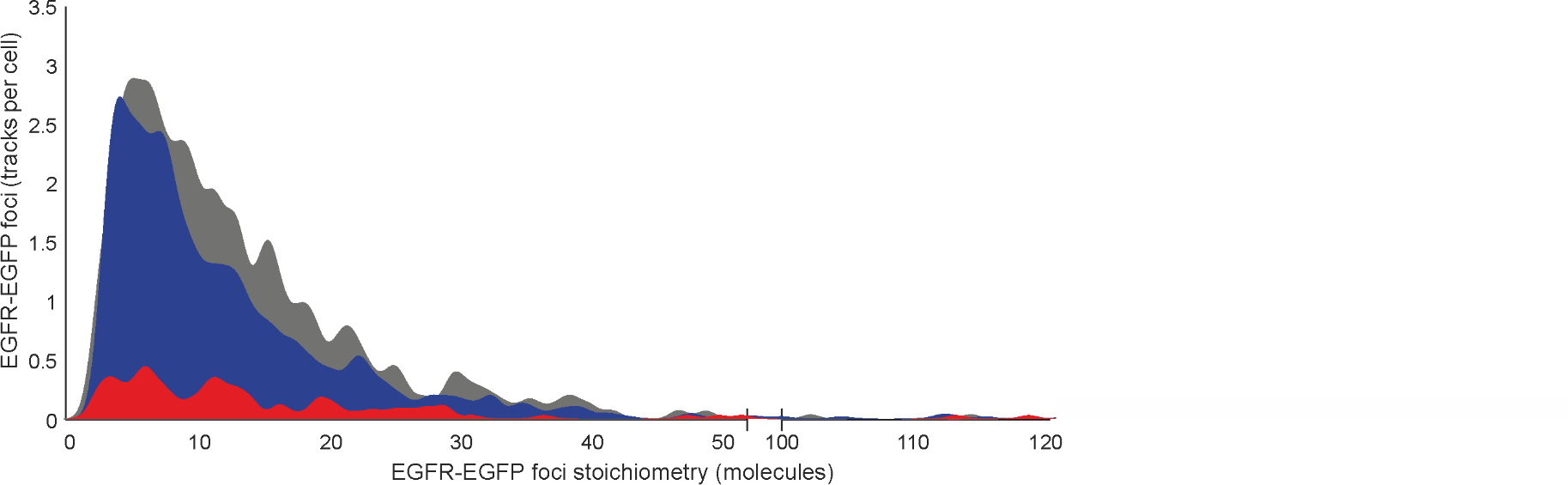

**Fig. S9. Effect of pertuzumab on EGFR foci stoichiometry.** Distribution of EGFR foci stoichiometry for cells treated with pertuzumab or, showing pre (grey) and post EGF addition for EGF-EGFR (red) and unligated EGFR (blue) foci, data collated across 60min EGF incubation time. Number of cells per dataset in the range N =21.

**Supplementary movie legends**

**Movie S1.** Live transected SW620 cell single-color TIRF imaging. TIRF movie showing two adjacent cells transfected with EGFR-GFP (green) before addition of EGF.

**Movie S2.** Live transected SW620 cell dual-color TIRF imaging. TIRF movie showing a single cell transfected with EGFR-GFP (green), 10min post addition of EGF-TMR (red, 100 ng/ml).

**Movie S3.** Live CHO-K1 cell dual-color TIRF imaging. TIRF movie single CHO-K1 cell transfected with GFP labelled HER2 (green) and EGFR labelled with HaloTag650 (magenta).

**Movie S4.** Live CHO-K1 cell dual-color TIRF imaging zoom-in. Zoom-in of cell shown in movie S3 indicating transient co-diffusion of HER2 and EGFR.

**Supplementary references**

45.	S. Sigismund, *et al.*, Clathrin-independent endocytosis of ubiquitinated cargos. *Proc. Natl. Acad. Sci. U. S. A.* **102**, 2760–2765 (2005).

46.	P. J. Enriori, *et al.*, Breast cyst fluids increase the proliferation of breast cell lines in correlation with their hormone and growth factor concentration. *Clin. Endocrinol. (Oxf).* **64**, 20–28 (2006).

47.	X. Michalet, Mean square displacement analysis of single-particle trajectories with localization error: Brownian motion in an isotropic medium. *Phys. Rev. E* **82**, 041914 (2010).

48.	K. G. N. Suzuki, *et al.*, Transient GPI-anchored protein homodimers are units for raft organization and function. *Nat. Chem. Biol.* **8**, 774–83 (2012).

49.	A. Kusumi, *et al.*, Paradigm shift of the plasma membrane concept from the two-dimensional continuum fluid to the partitioned fluid: High-speed single-molecule tracking of membrane molecules. *Annu. Rev. Biophys. Biomol. Struct.* **34**, 351–378 (2005).

50.	S. Felder, J. LaVin, A. Ullrich, J. Schlessinger, Kinetics of Binding, Endocytosis, and Recycling of EGF Receptor Mutants. *J. Cell Biol.* **117**, 203–212 (1992).

51.	M. R. Holbrook, J. B. O’Donnell, L. L. Slakey, D. J. Gross, Epidermal growth factor receptor internalization rate is regulated by negative charges near the SH2 binding site Tyr992. *Biochemistry* **38**, 9348–56 (1999).

52.	R. S. Kasai, *et al.*, Full characterization of GPCR monomer-dimer dynamic equilibrium by single molecule imaging. *J. Cell Biol.* **192**, 463–480 (2011).

53.	E. C. Hulme, M. A. Trevethick, Ligand binding assays at equilibrium: Validation and interpretation. *Br. J. Pharmacol.* **161**, 1219–1237 (2010).

54.	J. Barretina, *et al.*, The Cancer Cell Line Encyclopedia enables predictive modelling of anticancer drug sensitivity. *Nature* **483**, 603–607 (2012).

55.	M. C. Leake, D. Wilson, B. Bullard, R. M. Simmons, The elasticity of single kettin molecules using a two-bead laser-tweezers assay. *FEBS Lett.* **535**, 55–60 (2003).

56.	M. C. M. C. M. C. Leake, D. Wilson, M. Gautel, R. M. R. M. M. Simmons, The elasticity of single titin molecules using a two-bead optical tweezers assay. *Biophys. J.* **87**, 1112–35 (2004).

57.	S. R. Needham, *et al.*, Measuring EGFR separations on cells with ~10 nm resolution via fluorophore localization imaging with photobleaching. *PLoS One* **8**, e62331 (2013).
